# Estimating intracranial pressure via low-dimensional models: toward a practical tool for clinical decision support at multi-hour timescales

**DOI:** 10.1101/2020.06.26.174540

**Authors:** J.N. Stroh, T. Bennett, V. Kheyfets, D. Albers

## Abstract

Broad clinical application of non-invasive intracranial pressure (ICP) monitoring using computational models requires a method of modeling ICP on the basis of easily measured patient data such as radial or brachial arterial blood pressure (ABP). These models may be highly complex, rendering them too slow for clinical and operational use, or may rely on data that is not consistently available. Coupling these models to an upstream vasculature component model decreases data requirements. For the purposes of clinical decision support at multi-hour timescales, two natural choices for model development are to increase intracranial model complexity or to include feedback mechanisms between ICP and vascular model components. We compare the performance of these two approaches by evaluating model estimates against observed ICP in the case of a slow hypertensive event from a publically available dataset. The simpler model with bi-directional feedback requires minimal identifiability and is sufficiently accurate over these timescales, while a more complex is difficult and expensive to identify well enough to be accurate. Furthermore, the bi-directional simple model operates orders of magnitude faster than the more anatomically accurate model when driven by high-resolution ABP. It may also be configured to use lower resolution ABP summary data that is consistently clinically available. The simpler models are fast enough to support future developments such as patient-specific parametrization and assimilation of other clinical data streams which are illustrated during the case of a complex ICP regime for a different patient. We present model comparisons to highlight the advantages of the incorporated simple model and its possible predictive power with further optimization.

## 1 Introduction

Traumatic brain injury (TBI) is a major public health problem. Intracranial hypertension (ICH) is common after TBI and can cause secondary injury by decreasing local or global cerebral perfusion[4, 2]. Therefore, clinical management of ICH after TBI is an important element of improving patient outcome. Endogenous control of intracranial pressure (ICP) includes cerebral autoregulation (CA) mechanisms which, when functioning properly, seek to maintain cerebral blood flow (CBF) across a wide range of arterial blood pressure (ABP). The mechanism of CA is to regulate CBF via constriction and dilation of local arteries [21], although various other mechanisms are proposed (*cf*. [3] and references therein).TBI is often accompanied by elevated systemic ABP and a loss of cranial volume due to cerebral edema. Both of these reduce autonomic ICP regulation: the former may exceed the effective range of CA function, while the latter diminishes this range. The Monro-Kellie doctrine [39] postulates a constant volume of intracranial parenchyma (functional brain tissue) and fluids, so changes in net blood volume transport yield changes in ICP. Consequently, treatment of elevated ICP must also account for changes in systemic ABP, which is the external ICP driver under this hypothesis. Clinical protocols therefore seek to control ICP while maintaining cranial perfusion pressure (CPP, the difference between ABP and ICP) [28], or risk cerebral hypoxia.

Important changes in patient ICP occur at minute-to-hour timescales and clinicians need to know about them quickly. Decisions regarding escalation of care for TBI patients are often driven by elevated ICP, typically defined as exceeding 20 mm Hg (1 mm Hg ≈ 133.3 Pa) [32]. This underscores the need to monitor ICP and identify critical changes. Importantly, this form of clinical decision support will need to predict ICP on timescales on the order of minutes-to-hours rather than seconds. Timescales only seconds-long would not provide enough warning for clinicians to intervene.

### The need for ICP estimation

ICP is monitored *in situ* either using an external ventricular drain (gold-standard) or a fiberoptic intraparenchymal catheter. Placement of an ICP monitor is an invasive procedure and exposes patients to additional risks such as infection and hemorrhage [34] which may adversely affect outcome. In some patients, the risks associated with this monitoring method are outweighted by the benefit of ICP- and CPP-guided therapy, but patient selection is critical. Non-invasive ICP (nICP) estimation is less risky than invasive monitoring and could inform patient selection and ICP monitor placement timing (e.g. early for those who are predicted to benefit). In addition, nICP forecasting paired with invasive ICP monitoring could be a powerful anticipatory clinical decision support tool.

Generally, nICP estimation involves identifying a relationship between ICP and proxies that may be more easily observable in real-time. Such relationships may be explored empirically or on the basis of explicit models representing underlying physiology; a recent comprehensive survey of nICP estimation modalities is available [17].

### Data and clinical availability

Estimation of ICP using models and/or proxy data is highly dependent on the availability of specific data, which limits its usage. For example, nICP may be statistically estimated from concurrent measurements of ABP and CBF velocity from empirical relationships [30] or via physiological parameters [16, 9]. This velocity data is typically observed via transcranial doppler sonography and is limited by joint availability of the sonography apparatus and a trained instrument technician to properly localize observations to the the middle cerebral artery (MCA). Such data must then be available in a timely manner at sufficiently high resolution for quality control before use in clinical nICP estimation. CBF may also be estimated indirectly by near-infrared spectroscopic analysis of cerebral oxygenation which indicates the level of CPP [18]. While nICP may be estimated using a number of different modalities, practical considerations such as availability of data and clinical logistics render their applications difficult.

### TBI modeling and decision support

Mathematical models built upon additional physiological components can circumvent the strong data requirements by coupling nICP estimation to an upstream hemodynamic model. For example, the autoregulatory electrical analog model of [13, 29] is coupled to a hemodynamic model of major vessels above the aorta. This approach allows for continued refinement and augmentation of the model by coupling additional components to increase physiological fidelity at the expense of computational overhead. Highly complex mathematical models such as the low-dimensional whole-body physiological presentation of [20] include lymphatic and venous circulation mechanisms. However, such a model is focused on systemic dynamics and may be too coarse or slow for the purpose of clinical nICP estimation.

The current nICP estimation models considered here are not designed with practical, universal applicability, nor hours-long simulations in mind. Bridging the gap between current models and clinical need requires that the former be fast enough to produce clinical decision support at timescales relevant for the latter using commonly available data. The more anatomically-representative model of [29] estimates nICP from ABP without additional data, but emphasizes pulse-scale pressure signals rather than hour-scale dynamics. The fast nICP estimation schemes of [9] track ICP at suitable multi-hour timescales, but have stringent requirements for uncommon data which limits applicability. These studies (*viz*. [29, 37] and [16, 9]) and the models developed therein are cited extensively in this document; although contrapuntal to one another, they are both foundational to this study.

### Objectives of the paper

The perspective of this work is that ideal clinical support tools for TBI management include a model which estimates multi-hour nICP from commonly available data using both internal systemic pressure feedback and intracranial (IC, as an adjective) process resolution. Such a model is not currently known. This investigation considers two natural first steps toward it: to strongly couple simple ICP estimation schemes to a hemodynamical model, or to use a complex ICP estimation model with more accurate representation of IC physiology and its local processes. We present advantages and disadvantages of each approach to better inform development toward a tool representative of the ideal model.

This paper has three primary objectives. The first is to extend the nICP estimation framework of [9] by using a coupled arterial vasculature model to eliminate its dependence on jointly-measured CBF. The second is to evaluate this model in relation to similar approaches for ABP-based nICP estimation over a duration of hours. The third goal is to motivate additional model machinery, such as case-specific parameter estimation and inference, needed to implement the proposed estimation framework for complex, clinically important situations. These goals aim to develop and validate a practical tool capable providing timely support in the clinical decision making process for TBI patients on a broader timescale than those considered in the literature. The remainder of this study is organized as follows. Section 2 presents the models and method of investigation, describes model experiments, and establishes model assessment criterion. Section 3 presents results of the experiments and compares the models, and discusses simulations of more complex patient injuries which are poorly simulated without optimization. Section 4 summarizes the analysis and motivates ongoing work toward modeling nICP estimation in a particular direction on the basis of results and implications.

## 2 Materials and methods

The comparison of nICP estimation schemes the involves three essential parts – model configurations, aortic inflow data which drive the system, and metrics used to compare models on the basis various aspects of performance – which are presented in the following sub-sections.

### 2.1 Numerical nICP estimation frameworks

The models considered here are algorithms which transform aortic ABP data into nICP estimates using two components which may be coupled or independent. The first component is a vascular hemodynamics model which distributes ABP forcing through the systemic arterial network (AN) to the CoW, and is referred to as the AN-CoW. The second component, referred to as the intracranial model (ICM), estimates nICP estimates using outflow of the AN-CoW at cranial arteries. We evaluated ICMs that either consider the brain as a single compartment or as 6 compartments defined by the distributions of the anterior, middle, and posterior cerebral arteries. Considered model formulations are differentiated by whether they interact uni- or bi-directionally with the arterial network and by the complexity of the ICM component. The following configurations are possible and illustrated in Fig 1:

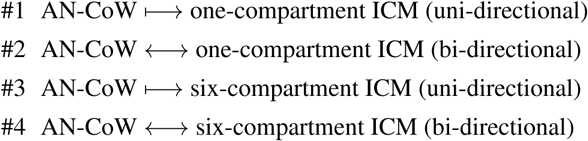

**Figure 1:**
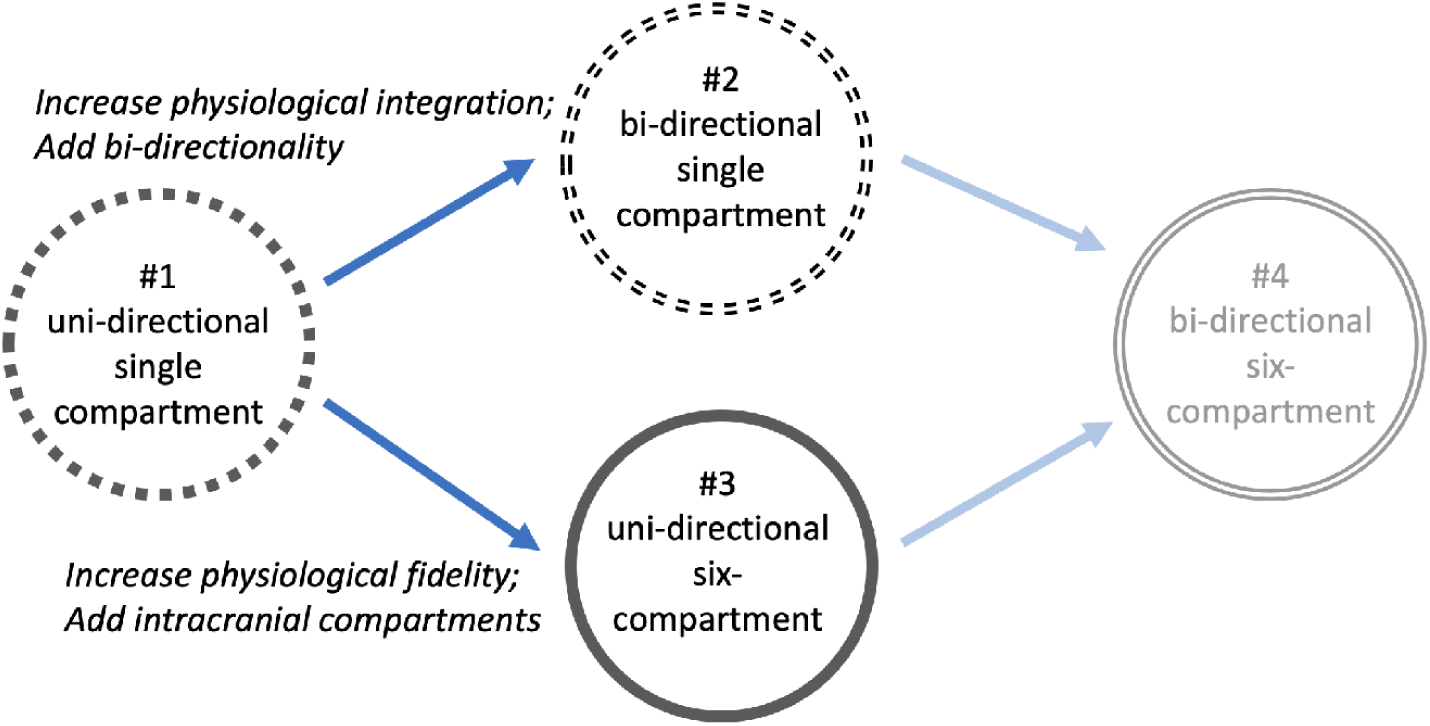
Conceptual overview of the relation between four models. The single-compartment model forced by prescribed ABP/CBF is the baseline model for comparison [16, 9] and is labeled as model #1. Model #2 integrates the lower arterial network forcing and single-component ICM into a common system through bi-directional coupling. In contrast, the six-compartment ICM [29] increases the physiological fidelity, model complexity, and parametrization of intracranial processes. Model #3 identifies this multi-compartment ICM uni-directionally coupled with the arterial forcing network. Model #4 is representative of the multi-compartment ICM fully integrated with the systemic arteries [29].

In the uni-directional configurations, the AN-CoW boundary outflow at the middle cerebral artery (MCA) is prescribed to the ICM as an inflow boundary condition. The AN-CoW calculates this pressure and flow for the entire simulation, which is then applied to the ICM. Bi-directional coupling of the AN-CoW and ICM enforces interactive agreement of flow volumes and pressures at the components’ interface (enforced as voltage and current electrical conservation).

Two directions for refining the base model are proposed as possible steps toward achieving a preferred but demanding model. Fig 1 shows the relationships of the models using model #1 as the most basic form, with models #2 and #3 as parallel steps toward ideal model #4. Models #2 and #3 extend model #1 either by a bidirectionally-coupled interface between the AN-CoW and ICM or by increasing the physiological complexity of the ICM component. This approach also tests which choice yields the highest gain in improvement over model #1 and the the cost of implementing it. Model #4 reflects an ultimate goal of a fully-integrated bi-directional model featuring an anatomically accurate ICM. However, such a model is not presented here due to its difficult implementation and impractical computational cost for the simulation timescales considered. Bi-directional coupling is difficult for multi-compartment models due to co-dependency of ICM state and the common pressure at each CoW terminal interface. Solving the nonlinear ICM state at each timestep takes several iterations, and each of these iterations requires recalculation of the entire upstream AN-CoW system constrained by pressure equality among the interfaces. The modeling framework used in this study is shown in Fig 2.

**Figure 2:**
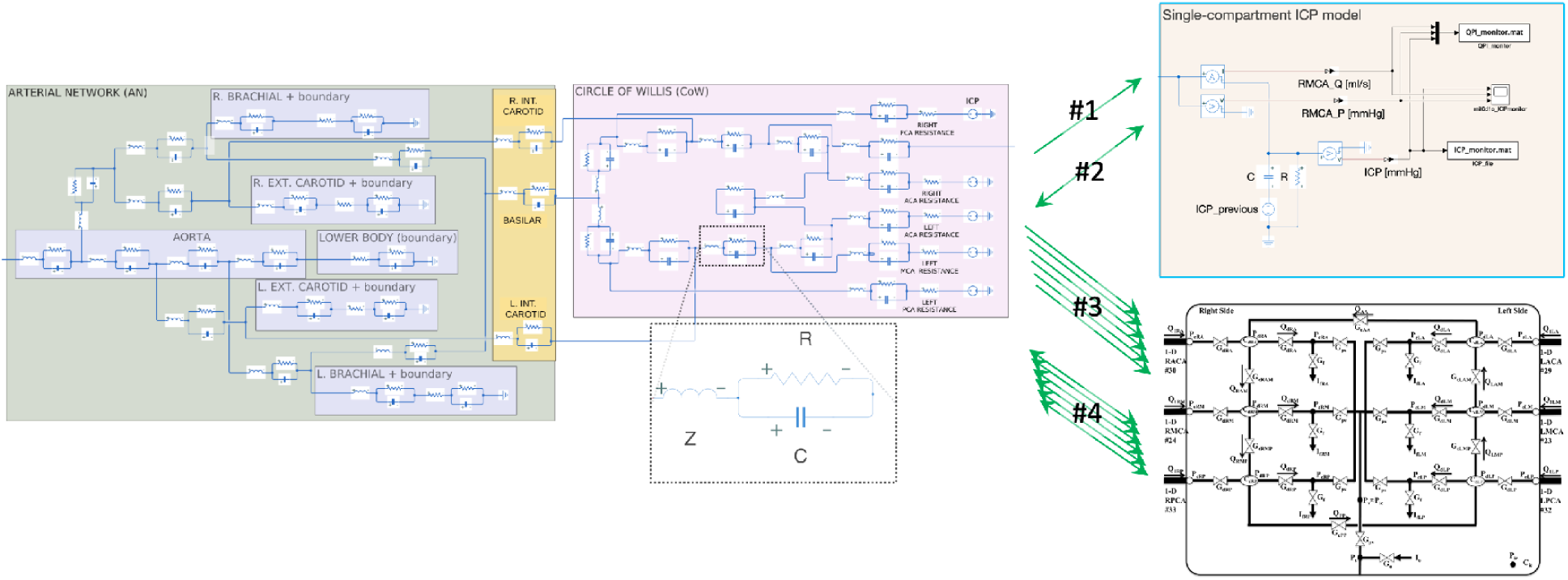
Diagram of model configurations 1–4. Schematic view of the various model configurations where green and pink boxes identify the AN and CoW model components, respectively, and ICMs at right. The aortic inflow pressure boundary condition is the sole driver of the system. Purple and orange boxes in the AN identify represented anatomy for reference. The AN-CoW is structured as in [29], but uses the 3-element (ZRC) electrical representations of vessels shown in the dashed white box. The single-compartment ICM is shown in the tan box; below it is the illustration of the 6-compartment model from [29]. Configurations #1–4 are distinguished by interactivity of coupling between AN-CoW and ICM, and by the type of ICM used; these are indicated by uni- and bi-directional green arrows between the model components.

Each model comprises two separate model components, which are described below. The AN-CoW for resolving hemodynamics outside the cerebral territories is presented in 2.1.1, while the ICMs for estimating ICP are presented in 2.1.2.

#### 2.1.1 Hemodynamical modeling of sub-cranial arteries

The AN-CoW model component is responsible for transforming aortic ABP data into flow and pressure at the MCA suitable for ICM input, and spatial resolution is unnecessary. Extra-cranial modeling of blood flow is a crucial component in nICP estimation models when patient ABP data are representative of lower vasculature states. Only terminal interface pressure and associated blood flow are required in this application. Therefore, the AN-CoW is modeled by a zero-dimensional framework of electrical analogs [15, 38], which are consistent with the physical equations [23]. This so-called lumped parameter approach has several advantges including a relatively small number of patient-specific parameters for each vessel and computationally efficient handling of vessel junctions and bifurcations. Further, conservation laws reduce at each timestep to algebraic systems rather than high-dimensional nonlinear functional representations [25] when spatially resolved.

The vascular system model is common to the various model configurations, and is shown schematically within Fig 2. The AN-CoW model comprised of a subcranial arterial network (green box) and the CoW vessels (pink box) are represented using 0D 3-element electrical analogs (white inset box) in MatLab SimuLink. Within it, vessel state variables pressure *P* and flow *Q* evolve (as voltage and current, respectively) under the influence of local vessel parameters (*R, C, Z*) and states of adjoining vessels. Base values for all vessel-level parameters and boundary conditions were adopted from previous studies (*viz*. [29] and references therein). Explicit definitions of vessel parameters and their relation to physical qualities are provided in Zero-dimensional vessel parametrization. Boundary conditions representing unresolved downstream vasculature are 3-element Windkessel models. To define AN-CoW outflow boundary conditions, fixed terminal resistances are set symmetrically for the ACA, MCA, and PCA such that flows target 1.3 ml/s, 2.2 ml/s, and 1.15 ml/s respectively, as in [29] for adults. In bi-directionally coupled models, CoW terminal vessels connect directly with the necessary IC vessels and require ICM pressure and resistance to coordinate with AN-CoW outflow. Specifically, the currently known estimates of pressure and resistance within the ICM are applied to the bi-directionally coupled MCA. Remaining uncoupled CoW termini are set as in the uni-directionally coupled case.

The large number of model parameters for the AN-CoW is reduced by considering a uniform scaling parametrization. The model of 33 vessels appears to have more than 100 total parameters in the RCZ framework, but each of these values are functions of anatomical dimensions length *l* and radius *r*. We assumed that vessel dimensions (*l, r*) scale uniformly within the AN and globally parametrized RCZ values according to proportionalities (*θ*_*l*_, *θ*_*r*_) in relation to the base values. This defines a nonlinear transformation of the electrical parameters via 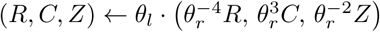. The remaining parameters – those for 3-element windkessel boundaries and CoW outflow resistance – are handled analogously with scalng parameters (*ω*_*l*_, *ω*_*r*_) and *R*_*term*_, respectively. Because CoW and adjacent vessel radii are approximately adult-sized by about 5 years of age [12], we did not scale vessels within the CoW model component.

Scaling the reference values *en masse* is effective within a realistic range of parameter values, as shown in Fig 3. This figure summarizes the relative effect of scaling parameters on properties of ICM inflow signals, determined by 500 simulations of parameters uniformly sampled from 0.5–1.5 for lengths and resistance and 0.9–1.1 for radii. Properties of ICM inflow signals are most sensitive to scaling of AN vessel dimensions and are less sensitive to scaling of the terminal resistance and AN boundary windkessel values. Scaling of AN vessel dimensions is more influential on the ICM than those related to CoW terminal resistance and AN boundary windkessel values. Further details are supplied in supplemental Fig 9. This re-parametrization reduces the AN-CoW component identification to five proportionalities (*θ*_*l*_, *θ*_*r*_, *ω*_*l*_, *ω*_*r*_, *R*_*term*_). It establishes a simple system-wide control over the vascular properties, improves parameter sensitivity, and provides a meaningful path to accurate model identification.

**Figure 3:**
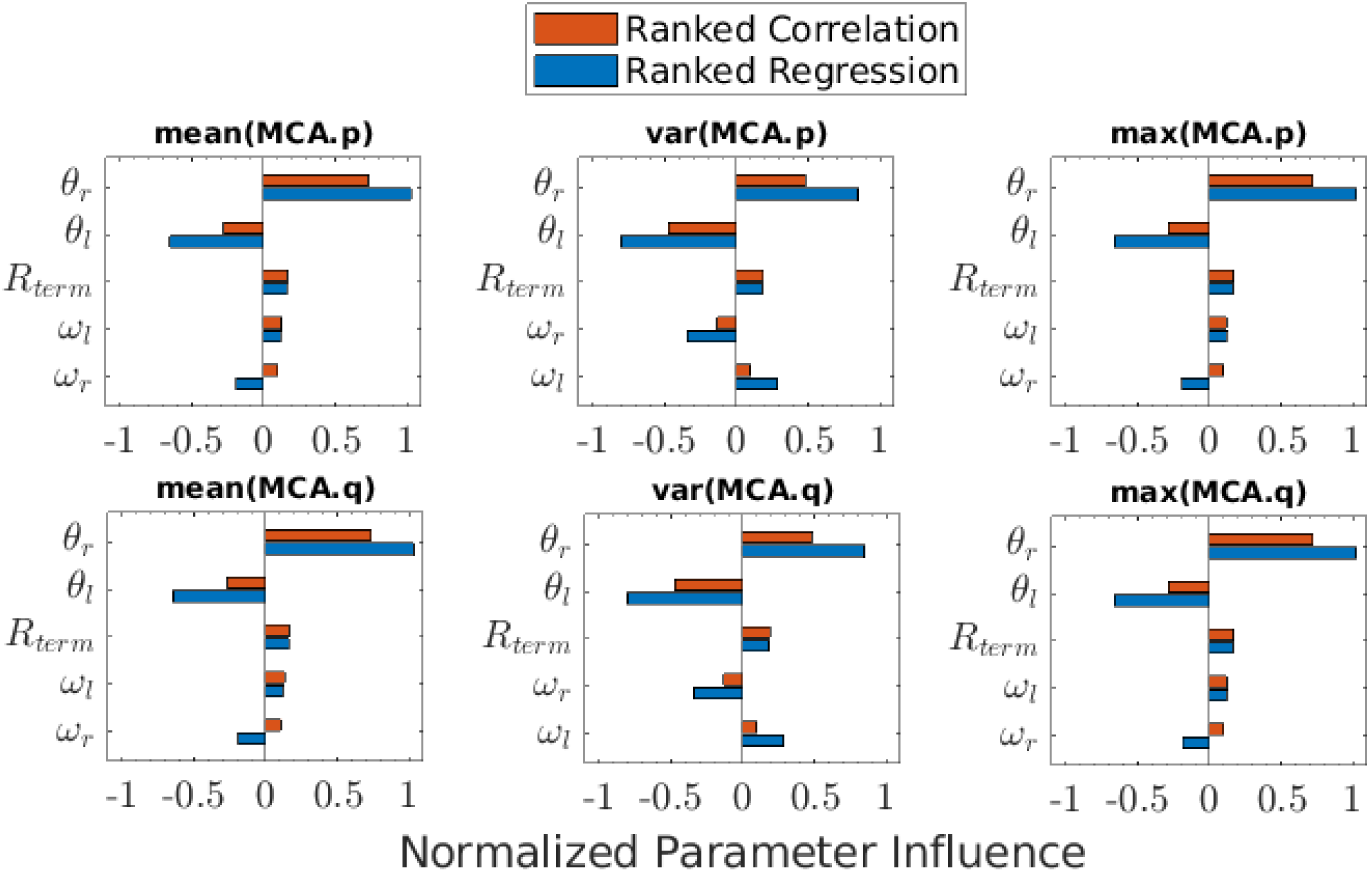
Ranked sensitivities of AN scaling parameters. Empirical estimates of sensitivity ranking, shown here for key signal features (mean, variance, and maximum) of MCA pressure (top row) and flow (bottom row), summarize Monte Carlo experiments using global structured random variations of scaling parameters (vertical axis of each panel) drawn from uniform distributions. Scaling parameters for the AN are most influential, while variations of terminal resistance and windkessel scale had a relatively little impact on solutions. Both regression rank and correlation rank are shown in normalized form. The regression predicts the linear change in MCA signal property with respect to change in a parameter among all changes in parameters, while the correlation measures the strength of the linear relationship between pairwise changes in parameter values and changes in MCA signal property. Note that rank ordering of windkessel dimensions is reversed for variance in MCA signal (center column). One concludes that scaling of vessel dimension parameters has considerable influence on ICM inflow signals, and provides global control while reducing the number of parameters needed to specify a patient-specific AN.

**Figure 4:**
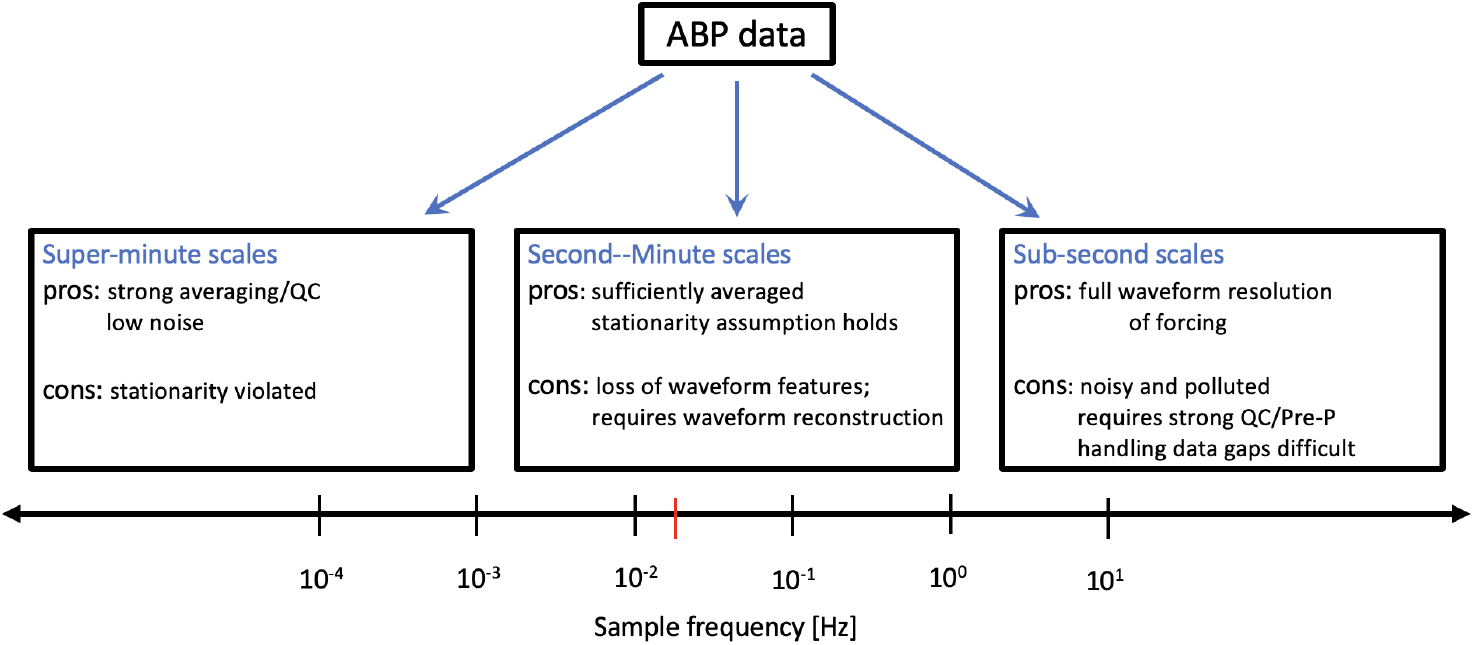
Timescales of ABP inflow data. The q1m data sampling frequency is indicated in red. Previous studies by [16, 9] validate the regressive method of the simple ICM using data sampled at 20–70 Hz and 125 Hz. The central scale is desirable for hour-scale applications as this resolution both qualitatively minimizes computational overhead and satisfies parameter stationarity. The latter assumption is necessary for the statistical schemes of the single-compartment models to function properly. The left-most scale offers strong smoothing and low noise, but fails to resolve pulsatile waveform and violates assumptions of the simple models.

**Figure 5:**
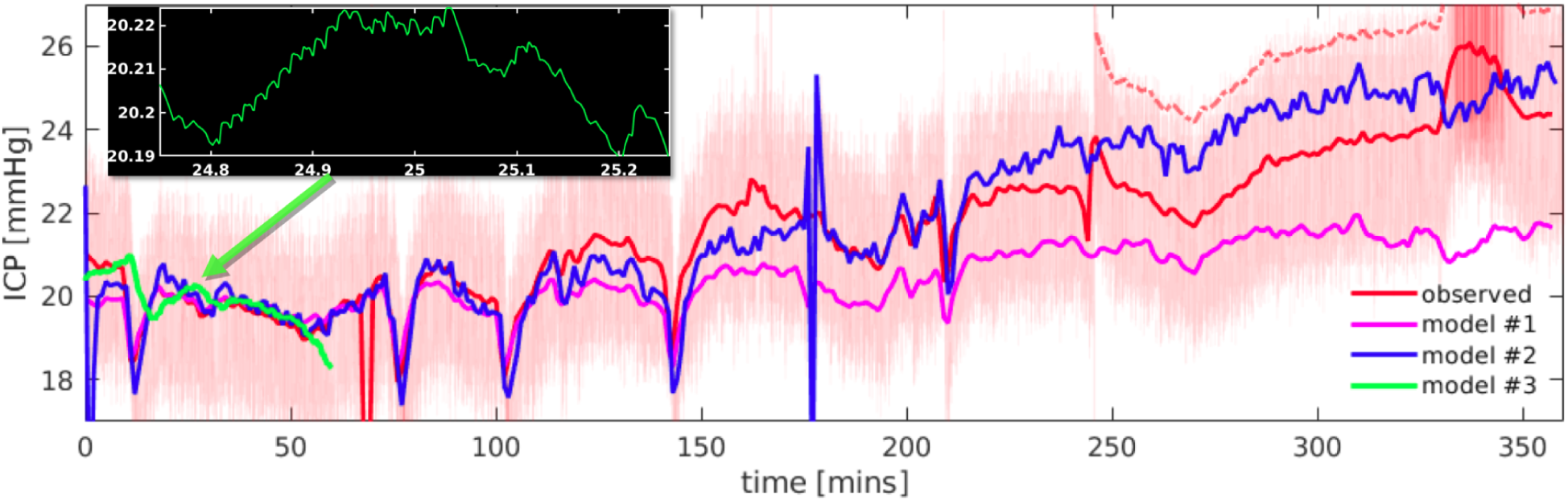
Observed and estimated nICP for patient #6. Depicted are observed (red) and estimated nICP for Charis patient #6 using models #1–3. Model #1 (magenta) takes approximately 14 minutes to run from offline-supplied MCA flow and pressure estimates; it has a bias of 6.6 mm Hg over the first hour. Model #2 (blue) takes approximately 34 minutes to run from ABP forcing including simulation of the AN-CoW; it has a bias of 6.4 mm Hg over the first hour. Model #3 (green) simulates nICP over 1 hour and takes approximately six hours of clock time; it has a bias of ∼6.4 mm Hg and requires a variance inflation scaling of 27 to obtain the curve shown. The black inset shows a 30-second interval of model #3 nICP to illustrate pulse resolution. After initial adjustment of baseline pressure, model #2 outperforms the other nICP estimates. Model #1 fails to follow the longer-term trend of rising ICP whereas #2 does, suggesting that bi-directional interaction between components is crucial. The large transient error in model #2 around 180 minutes results from errors in the ABP inflow.

**Figure 6:**
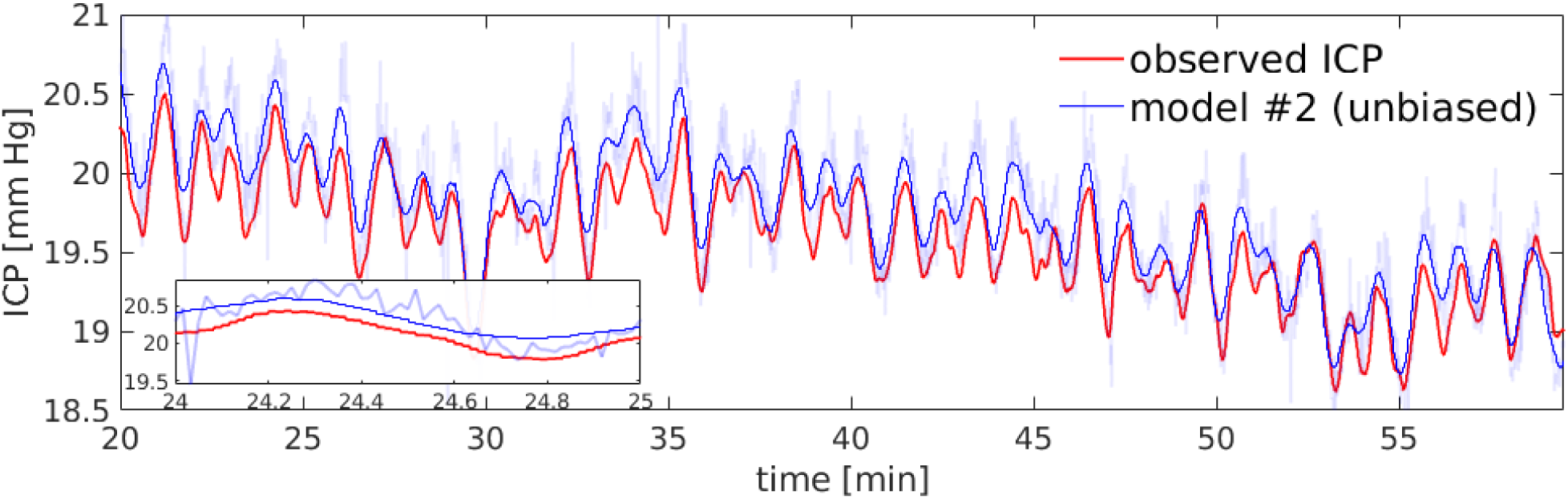
Strong local tracking of ICP signal in model #2 at the expense of computational time. The mean nICP estimates over 30 second intervals (blue curve) using the output of model #2 (light blue) with raw ABP strongly track observed ICP (red curve). The modeled simulation includes accurate reproduction of local trends and *O*(10^−2^) Hz waves of the averaged observed ICP. This simulation calculated resistance and compliance parameters at 1 second intervals using 30-second regression intervals (*i*.*e*. with 29 second overlap). The corresponding mean ICPs over those are plotted as solid curves for comparison with the observed ICP. While four times slower than real time, this simulation is roughly twice as fast as model #3 under pulsatile aortic inflow and requires no additional data or external inference.

**Figure 7:**
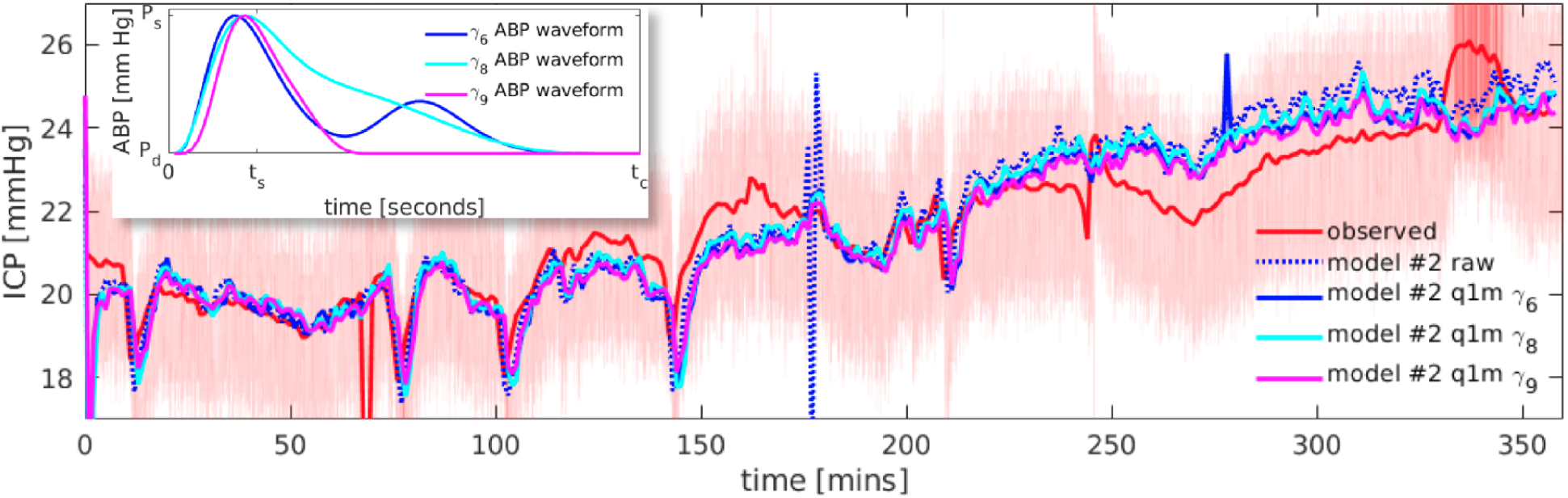
Model #2 performance using q1m summary ABP data. Various simulations under q1m inflow data are compared with observed ICP (red curve) and nICP estimate based on raw 50 Hz data (dashed blue). The nICP estimates using the correct q1m ABP data with both correct (blue) and incorrect (magenta and cyan) waveform parameters are also shown. Q1m simulations use ABP waveforms formed from q1m summary data projected onto continuous waveform versions with constant systolic and diastolic peak pressure and heart rate. In the inset figure: solid blue, cyan, and magenta lines show ABP waveform shapes for patients #6, #8, and #9, respectively. The nICP, plotted in corresponding colors, based on those waveforms track ICP well and are qualitatively indistinguishable. This shows that q1m ABP is sufficient for the aortic inflow and that patient-specific parametrization of ABP waveforms has little advantage.

**Figure 8:**
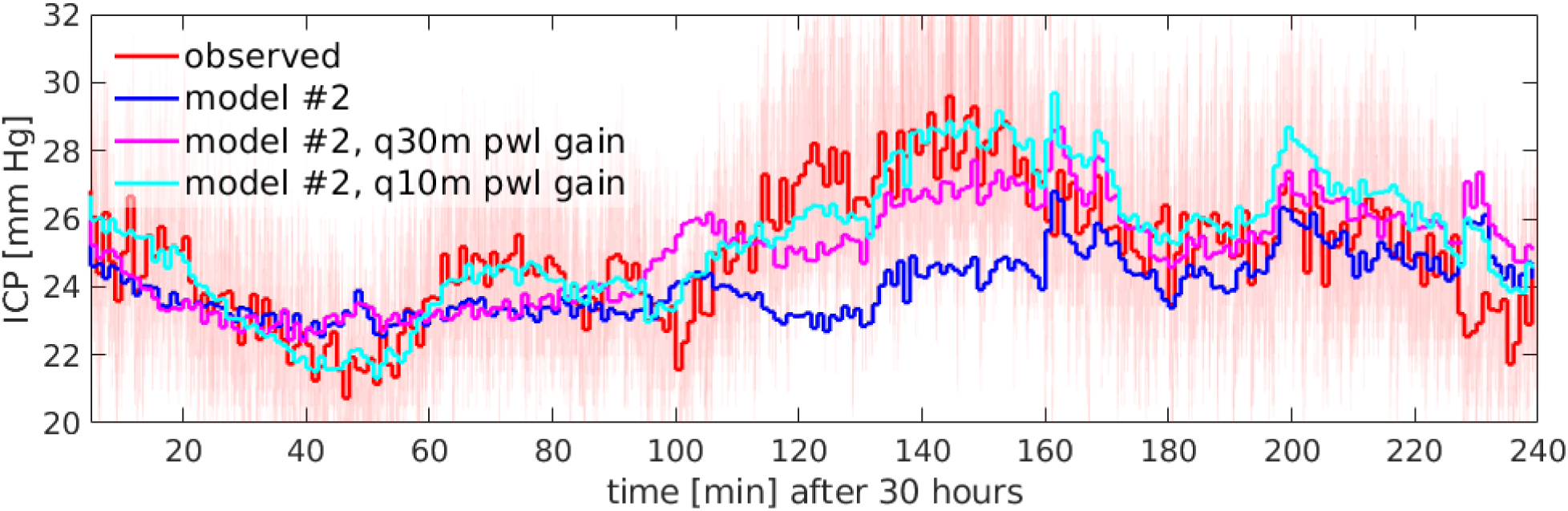
An illstrated complex case application: patient #5. The observed ICP signal for Charis patient #5 during record hours 30–34 is shown in red, with the dark red line indicating q1m average ICP as in previous figures. Signal noise and high-frequency variability are much stronger than in the records of patient #6, yielding a less smooth observed mean ICP. Slow wave pressure dynamics are observed during this period, but they are absent from the model #2 solution using optimized scaling parameters alone (blue curve). This nICP estimate is relatively constant with small variance (∼1 mm Hg) until 160 minutes into the simulation. The estimate also misses the onset of the ∼7 mm Hg ICH event during 100–140 minutes and begins to track it only after ICP begins to fall. Model experiments using additional non-stationary gain applied to inflow show improved trend tracking during these more dynamic regimes. The nICP estimate is improved greatly using gain parameters specified at 30 minute intervals (magenta curve), resulting from the addition of eight independent parameters. The large pressure event 40 minutes sooner in this simulation, but still fails to capture shorter-term ICP dynamics around 20–80 minutes. A similar gain specified at 10 minute intervals greatly improves the nICP estimation (cyan curve, using 24 parameters rather than 8) of the ICH event along with other smaller features including initial waves in the first 100 minutes.

**Figure 9:**
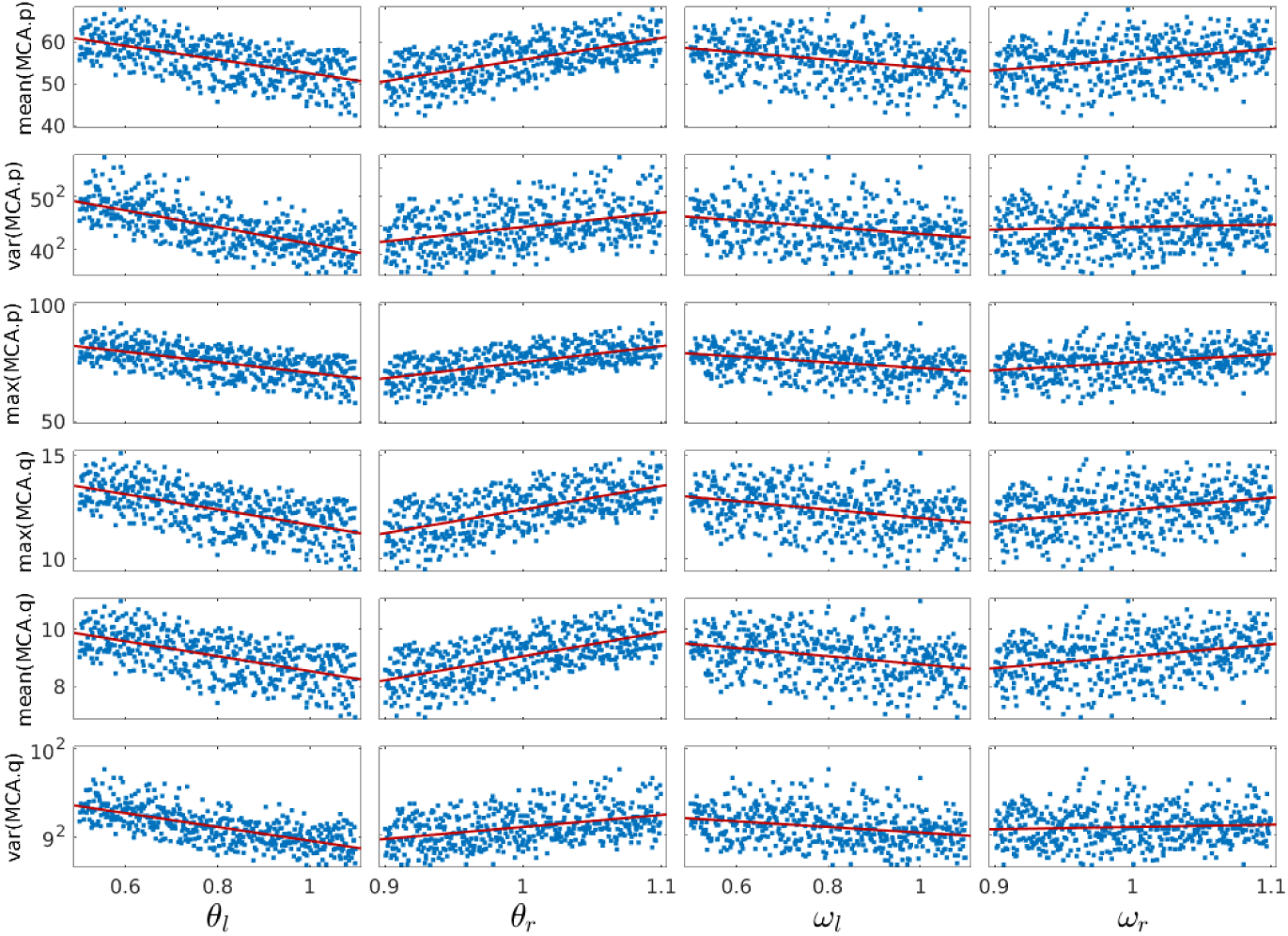
Monte Carlo sensitivity experiments. Monte Carlo sensitivity experiments performed on the base AN-CoW components with fixed ICP and IC parameters shows how scaling parameters (in columns, at bottom) affect IC component boundary forcing. Each 1-minute simulation (blue points) used artificial ABP forcing in the form of a 1Hz cycle comprised of a 0.15 second sinusoidal systolic upswing to 125 mm Hg followed by a 0.15 second return to 80 mm Hg diastole. The top three rows correspond to values of the mean, variance, and maximum of MCA pressure, and the bottom rows correspond to MCA flow. Columns, left to right, correspond to scale parameters for vessel length, vessel radius, windkessel length, and windkessel radius. Red lines establish the relationship between changes in scale parameters and the response in MCA signal properties; their slopes are used in the regression ranking of Figure 3. Random joint variations of each scale parameter (and terminal resistance scaling, not shown) sampled and assigned to 500 simulations via Latin hypercube sampling. Uniform sampling distribution ranges were [0.5, 1.1] for lengths, [0.9, 1.1] for radii, and [0.5, 2] for resistance, with weak and positive covariances (0.5) assumed between lengths (and radii) in an attempt to preserve anatomical fidelity.

#### 2.1.2 Intracranial Pressure model components

The ICM component is the algorithm responsible for estimating nICP from the AN-CoW outflow to cerebral arteries. The two ICM configurations considered are a six-compartment model based on [29] and [13] and a single-compartment model based on [16] and [9]. In addition to the number of represented cerebral perfusion territories, the models differ in their estimation approach. The multi-compartment model is more anatomically accurate and explicitly resolves IC hemodynamics with communicating arteries and autonomic pressure regulatory processes. The single-compartment approach, in contrast, computes ICP using window-based statistical estimates of IC compliance and pressure determined through regression of ICM inflow waveform properties. Overview of the multi- and single-compartment ICMs occur below under subheadings 2.1.2 and 2.1.2, respectively, with further technical details provided in supplements Six-Compartment ICM details and Single-compartment.

##### Overview of the six-compartment model

The complex model of [13, 29] presents an anatomical layout of the main cerebral pathways and their dependent mechanisms. Using six interacting territories, it includes IC pressure and perfusion dynamics coupled by communicating arteries, CA processes, and cerebrospinal fluid (CSF) balance. CA processes are modeled by internal feedback mechanisms which regulate flow through each compartment via vaso-constriction/-dilation [36]. This autonomic control influences the local pressure and flow balances between compartments, leading to inter-compartmental blood flow via communicating arteries. IC pressure and compliance are non-linearly co-determined by total IC volume changes resulting from CA processes and net fluid (blood + CSF) change. A mathematical description, including a list of physiological and model parameters, is provided in Six-Compartment ICM details. The high degree of physiological fidelity resolves IC dynamics at timescales inherited from ABP forcing, including the pulsatile ICP waveform [29]. Further, the 6-compartment nonlinear nICP component calculates numerous potentially clinically relevant diagnostic variables (List of diagnostic variables in the six-compartment models) during simulation.

##### Overview of the single-compartment model

The single-compartment ICM of [16] is a simple model which estimates ICP physiologically rather than modeling it anatomically. Here, nICP is constructed from least-squares estimates of bulk IC compliance (*C*) and resistance (*R*) over a temporal window containing several cardiac cycles. The algorithm follows [16] and [9], and is presented supplementally in Single-compartment. The underlying assumption is that ICM parameters are stationary throughout the estimation window. Succinctly, the model first estimates compliance *C* as a scaling factor between total MCA inflow volume and the corresponding pressure change during the systolic upswing of each pulse. The estimated compliance identifies blood flow (*Q*_1_) through a yet unknown resistance *R*. The value of *R* is subsequently estimated as the proportion of the change in applied pressure to associated change in flow *Q*_1_ through the unknown resistance. An nICP may be calculated algebraically as the difference between inflow pressure and the associated pressure lost by flow across the estimated resistance. However, the implementation here defines nICP as the mean simulated ICP forecast based on the values of resistance *R* and compliance *C* in the previous window.

The estimation process of this ICM requires no physiological parameters, but does require algorithmic parameters that influence model behavior. Two required model hyper-parameters control the length of the temporal window over which each estimation occurs and the timestep of parameter updates. The first is limited by the stationarity assumption and determines the sample size for the regressions, while the second controls output temporal resolution and coupling strength. In models #1 and #2, estimates of IC resistance and compliance inform the simulation of the next regressive window. However, the calculated nICP alters the outflow pressure condition at each CoW boundary when bi-directionally coupled (model #2). The length of the update timestep therefore affects the temporal coarseness of the nICP estimate in each model and also defines the timescale of feedback between the ICM and upstream vascular model in the bidirectional one. Single-compartment model simulations use 1 minute windows and 1 minute updates unless otherwise specified.

### 2.2 Observational Data and Patient Selection

The models require aortic inflow boundary data (referred to simply as ‘forcing’) pressure (or flow) measurements for several hours’ duration and a concurrently observed ICP for evaluation of model output. Additionally, models #1 and #2 require that boundary ABP be pulse-resolving. A suitable dataset for this purpose is the CHARIS v1.0.0 collection (‘Charis’ hereafter, [19]), which is publicly available online at PhysioNet [10]. These 50 Hz data comprise joint radial ABP and ICP timeseries of 13 patients with IC injuries. This study focuses on patient #6 of that data, a 20-year-old male with TBI, for model validation. This patient was selected for the simplicity of his injury, cleanliness of joint ABP-ICP signal, and representativeness of base parameters (*e*.*g*. optimal scaling parameters for the AN-CoW were approximately 1); he could simulated out-of-the-box. Other patients in the dataset are significantly older or have multiple documented brain injuries (*e*.*g*. ischemic or hemorrhagic stroke). Also, large scale noise or corrupt signals are common in the records of the patients (*cf*. Fig 10); their ABP and/or ICP data could not be utilized contiguously for 4–6 hours periods without extensive and uncertain pre-processing of the available data.

**Figure 10:**
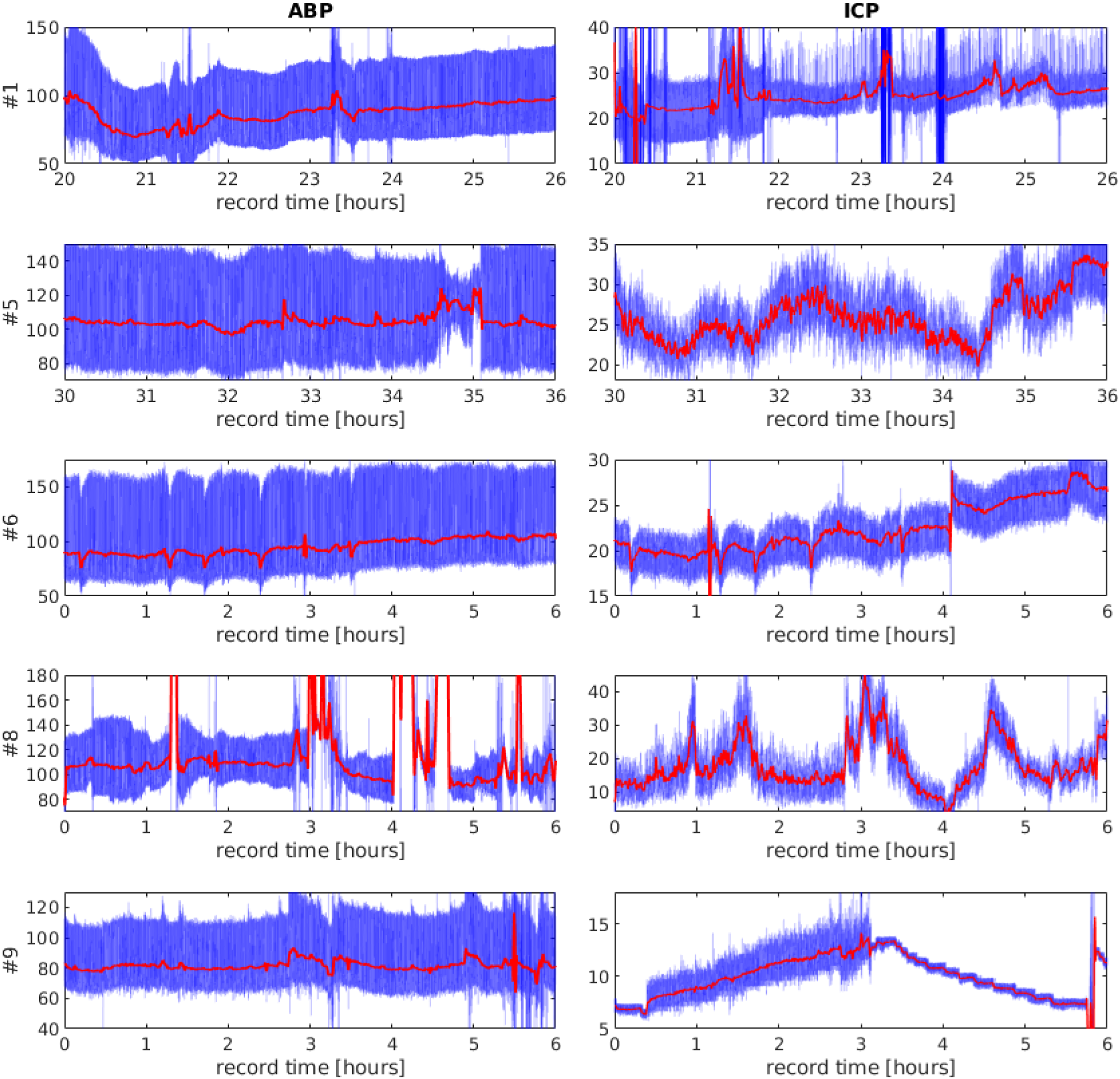
Selected Charis patient joint ABP-ICP timeseries. Fifty Hz ABP (left column) and associated ICP (right column) for Charis patients #1, #5, #6, #8, and #9 (rows top to bottom, respectively, labeled at left). Vertical axes corresponds to pressure measurements in mm Hg and horizontal axes show time in hours. Blue regions outline the raw pulsatile signal, with the red curve identifying the signal smoothed over 2 minute window by a cubic polynomial filter to illustrate the scale of local signal variability. Patient #6 was chosen for benchmarking on the basis of low noise in joint ABP-ICP signals and interpretability of ICP dynamics over a several-hour time period. Patient #5 was chosen for idealized optimizability experiments due the timescale of ICP dynamics continuity over a multi-hour interval, and the decorrelation between ABP and ICP during this period. Patients #8 and #9 are included here since their ABP waveform shapes strongly contrast those of patient #5 (*cf*. Figure 7 inset).

Using the Charis radial ABP data as aortic inflow in the models introduces errors which are consistent throughout all experiments. However, it was freely available and satisfied other aforementioned requirements. This usage is obviously incorrect and biases systolic pressure more than diastolic [27, 26].Sophisticated transformations exist [5] for reconstructing aortic pressure from radial ABP, but the simple approach taken here avoids uncertainties associated with that additional algorithmic processing.

The six-compartment models (#3, and presumptive #4) can act on many different types of aortic inflow conditions, including both raw and mean non-pulsatile forcing. Meanwhile, the simple regressive models depend strongly on pulsatile signals at the middle cerebral artery to identify periods of elastic vessel response and maximal resistive flow. Fig 4 identifies possible input data sampling frequencies. Given that systole is approximately 1/3 of the cardiac cycle (approximately 1 Hz), raw APB data samples should be at minimum 10 Hz to meet these criteria, even if downscaling is considered. On the other hand, data with a high sampling frequency is often noisy or inconsistent, and jointly measured signals may experience temporal decoupling through instrument clock drift. This implies that such datasets may require extensive preprocessing and/or resampling prior to use (*cf*. [9]).

An additional mechanism is proposed to define aortic inflow from lower-resolution ABP summary data. As outlined in the introduction, liberating nICP estimation methods from dependence on high-frequency measurements requires additional mathematical techniques to estimate them from more commonly available data. Further consideration is therefore given to ABP forcing in single-compartment models with the aim of using clinical data in continued development. Specifically, lower-frequency ABP records are assumed to comprise non-overlapping one-minute averages of systolic and diastolic pressures and heart rate (*sp,dp*, and *hr*, respectively). To this end, a representative portion of raw 50 Hz patient ABP was used to identify static, patient-specific waveform parameters *γ* for a function of the form

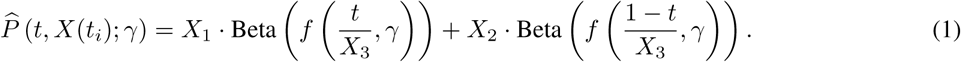

Here the symbol *X*(*t*_*i*_) = [*sp, dp, hr*]_*i*_ identifies the averaged ABP information during interval *i* of a *quaque* 1-minute (q1m) record, defining values for *t*_*i*_ <= *t* <= *t*_*i*+1_. The function 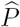 transforms q1m information into a piece-wise uniform patient-specific pulsatile signal at an arbitrary resolution. Inflow signals constructed thusly from Charis ABP data are conveniently sampled at 60Hz. In relation to Fig 4, this process maps q1m data (identified by the red mark) into the scale useable by models #1 and #2.

### 2.3 Measures of Quality and Efficiency for models

To evaluate each experiment, we calculated three scores for each nICP estimate based on accuracy, precision, and speed for simulations over time interval [0, *T*]. The symbol nICP* in this discussion indicates nICP debiased against observed ICP during the first hour of simulation. Justification for this correction is that skill scores evaluate model ability to track variability in recorded ICP data rather than estimate the absolute pressure. It also accounts for the bias induced through mis-use of radial ABP as aortic inflow pressure. Each evaluation is applied to an nICP estimate, the score of which is then associated to the model which produced it.

The first score

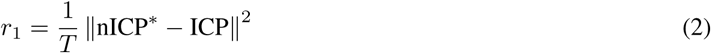

rates the ability of the model nICP to track changes in observed ICP. Normalization by simulation duration *T* is needed to compare experiments of different lengths. This score is the time-average standard error between ICP and model estimate, which quantifies how generally inaccurate a model nICP estimate is.

The second evaluation, with **⟦** · **⟧** denoting the logical test operator, is

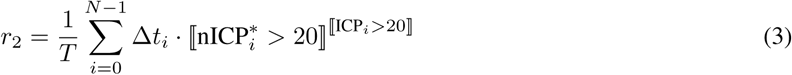

which is the mean percentage of simulation time that nICP correctly agrees with observed criticality (ICP>20 mm Hg) over timesteps 0 ≤ *i* ≤ *N*. Although seemingly qualitative, it is more clinically relevant than *r*_1_ as it quantifies whether nICP provides accurate and actionable information to support decision making.

Finally, the third quantity is simply

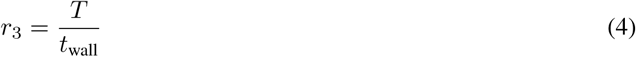

or the ratio of simulated time interval to elapsed clock time, with *r*_3_ *>* 1 indicating faster-than-real-time forward model integration. Values of *t*_wall_ correspond to serial run times calculated using Matlab R2020a on a 2019 model iMac Pro with a 3.7 GHz Intel i5 CPU. This final evaluation measures how practical a model is for providing timely clinical support as well as its utility in other applications, such as nonlinear parameter estimation or data assimilation applications which require extensive, repeated forward model integration.

The number of necessary parameters required for realistic initialization and the input data fidelity are among other aspects not evaluated quantitatively. They are discussed in the context of model utility, but not scored explicitly. Finally, all model simulations are initialized with zero-flow within the AN-CoW system. This component is common among the various model configurations and its algebraic nature distributes flow instantaneously. A brief spin-up adjustment period occurs during the first 2–3 minutes of simulation (2–3 ICM parameter updates in models #1 and #2) and these errors included in skill calculation with negligible impact on comparative assessment.

## 3 Results

### 3.1 Comparative assessment of model simulations

We ran model configurations #1–#3 for the first hours of patient #6 record data to evaluate nICP and efficiency on the basis of the skills presented above. Fig 5 shows the observed ICP signal along with estimates from each model. During the simulated period, patient ICP exhibits variability about ∼20mm for about 2.5 hours, followed by a gradual non-monotonic rise to ∼23 mm Hg. Sharp temporary decreases in ICP around 10mins, 75mins, 105mins, and 142mins probably result from interventions (mannitol or hyperventilation treatments) to reduce ICP [19]. The observed ICP signal used in model evaluation is plotted in solid red. A signal discontinuity near 243.83 minutes, where ICP data increases by 5+ mm Hg within one minute, may be due to transducer recalibration. To compensate, observed ICP is decreased by ∼2.5 mm Hg after 243.5 minutes. The original unaltered one-minute average ICP observations (dashed light red) are shown for reference over the interval 244–360 minutes.

Model comparison is organized below into three subtopics discussing qualitative differences, quantitative differences, and observations about resolvable timescales and fidelity.

#### 3.1.1 Qualitative differences between modeled nICP series

Models #1 and #2 produce qualitatively different nICP estimates, with the key difference being that model #2 follows the multi-hour trend of increasing ICP. Model #1 tracks the observations well until around 2 hours, but it fails to follow subsequent rise in observed ICP after the intervention around 142 minutes. The bias of model #1 over this interval is approximately −1.8 mm Hg and increases over time. In contrast, model #2 tracks this observed ICP rise, although it overestimates ICP by an average of 1.02 mm Hg during 220-360 minutes. Both models #1 and #2 also underestimate the ICP local maxima around 130 mins and 160 mins. One concludes that inclusion of bi-directional coupling improves estimation of low-frequency ICP signal components that are crucial in applications spanning several hours. Interestingly, neither model resolves the 2 mm Hg humplike event of the observed signal occurring during 330–350 minutes of the patient record. This feature in the observed ICP could result from a temporary change in patient posture, but no corresponding change occurs in the aortic ABP inflow signal (*cf*. 10, center left panel). This provides evidence that changes in ICP not arising from aortic ABP dynamics may not be resolved by simple ICMs.

Model #3 is simulated only for the first hour due to slow computation and inaccurate parameters which require additional inference to determine. This model is difficult to initialize correctly due to the number of parameters in the six-compartment ICM. Exploratory changes in ICM parameters while attempting to obtain realistic behavior often led to divergent nICP estimates during the first hour of simulation. This indicates a strong dependence on dynamically representative and balanced parameters that must be inferred *a priori* in a practical implementation. ICM parameters representing venous capillary conductance (*G*_*pv*_) and reference pressure (*P*_*icn*_) were tuned sufficiently to obtain the reported nICP estimate. This solution also includes mean variance inflation by an optimal factor 26.3 to account for remaining uncalibrated parameters. The modified solution estimates the observed trend well, although the localized variance is too small. It also lags the behind observed ICP by approximately 228 seconds. This apparent delay, like the reduced variance noted at several timescales, likely reflects poor representation by generic ICM parameters in the absence of additional inference. Further attempt to more accurately prescribe these parameters was obstructed by long runtimes, which were approximately six hours per simulated hour.

#### 3.1.2 Quantitative differences between modeled nICP series

The qualitative advantages of the bi-directional simple model over the uni-directional complex one are borne out by model skills *r*_1_–*r*_2_ shown in Table 1. Comparing scores reveals that model #2 has a practical advantage over model #3 in terms of identifiability and is able to easily and quickly simulate a six-hour period. Because the longer simulations of models #1 and #2, scores are calculated separately for the initial 1-hour period resolved by all three models and appear with parentheses. Bi-directional coupling improves simple model accuracy by nearly 30% and makes identifying critical ICP 12% more accurate over the 6 hour duration. Over the initial hour, there is a slight loss in accuracy due to longer spinup adjustment and a slight improvement in critical ICP identification. This further supports that the feedback mechanism improves low-frequency tracking which has no advantage over short timescales. Use of the uni-directionally coupled complex ICM in model #1 leads to a marginal (<1%) improvement in critical ICP identification over the base model, but incurs a 57% loss of analytical accuracy compared with the base model. Most of this loss is because of the ∼4-minute lagged response of the ICM. Accounting for it shows a considerable decrease in error to *r*_1_ = 2.75 accompanied by a loss in specificity to *r*_2_ = 0.84. Model ranking of model #2 over model #3 on the basis of skill is unchanged for the isolated first hour even with posterior modification of model #3 output.

**Table 1:** Model scores for principal comparison.

| | $r_1$ [accuracy] | $r_2$ [precision] | $r_3$ [speed] |
| --- | --- | --- | --- |
| Model #1 | 5.01(2.42) | 0.877(0.92) | **116.129** |
| Model #2 | **3.53**(2.47) | **0.97**(0.98) | 7.356 |
| Model #3 | (3.83) | (0.883) | (0.145) |

Scores for simulations of Charis patient #6 during initial hours of data, with best results for each score shown in bold. Scores $r_1$ and $r_2$ rate the nICP accuracy and ability to identify critical ICP, respectively, while score $r_3$ rates the speed of the nICP estimation process. Parenthesized entries are calculated using only the first simulated hour.

**Table 2:**
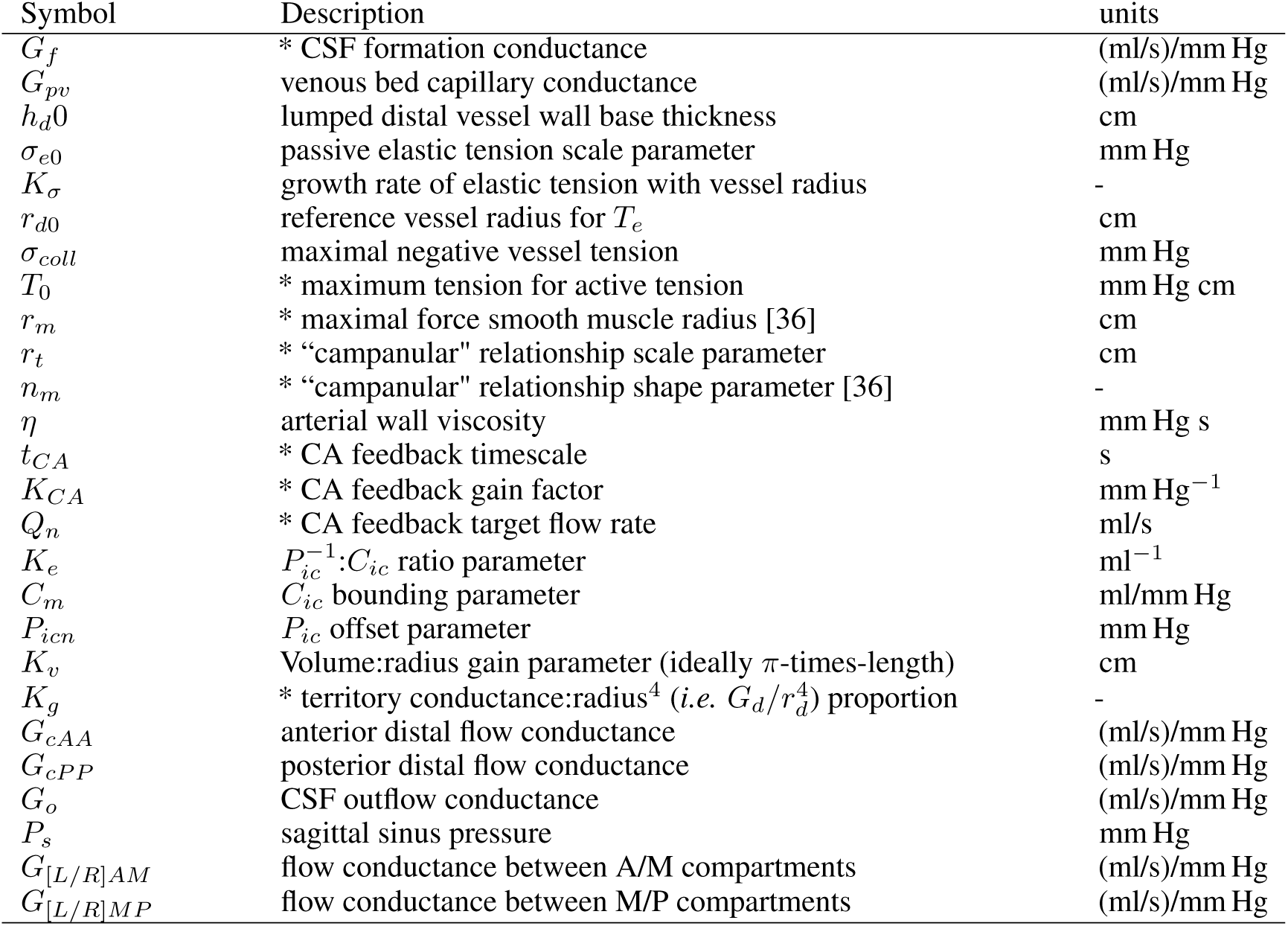
List of required primitive parameters in the six-compartment ICM. Parameters indicated by * may vary between each compartment.

| Symbol | Description | units |
| --- | --- | --- |
| $G_f$ | * CSF formation conductance | (ml/s)/mm Hg |
| $G_{pv}$ | venous bed capillary conductance | (ml/s)/mm Hg |
| $h_{d0}$ | lumped distal vessel wall base thickness | cm |
| $\sigma_{e0}$ | passive elastic tension scale parameter | mm Hg |
| $K_\sigma$ | growth rate of elastic tension with vessel radius | - |
| $r_{d0}$ | reference vessel radius for $T_e$ | cm |
| $\sigma_{coll}$ | maximal negative vessel tension | mm Hg |
| $T_0$ | * maximum tension for active tension | mm Hg cm |
| $r_m$ | * maximal force smooth muscle radius [36] | cm |
| $r_t$ | * "campanular" relationship scale parameter | cm |
| $n_m$ | * "campanular" relationship shape parameter [36] | - |
| $\eta$ | arterial wall viscosity | mm Hg s |
| $t_{CA}$ | * CA feedback timescale | s |
| $K_{CA}$ | * CA feedback gain factor | mm Hg<sup>-1</sup> |
| $Q_n$ | * CA feedback target flow rate | ml/s |
| $K_e$ | $P_{ic}^{-1} : C_{ic}$ ratio parameter | ml<sup>-1</sup> |
| $C_m$ | $C_{ic}$ bounding parameter | ml/mm Hg |
| $P_{icn}$ | $P_{ic}$ offset parameter | mm Hg |
| $K_v$ | Volume:radius gain parameter (ideally π-times-length) | cm |
| $K_g$ | * territory conductance:radius<sup>4</sup> (*i.e.* $G_d/r_d^4$) proportion | - |
| $G_{cAA}$ | anterior distal flow conductance | (ml/s)/mm Hg |
| $G_{cPP}$ | posterior distal flow conductance | (ml/s)/mm Hg |
| $G_o$ | CSF outflow conductance | (ml/s)/mm Hg |
| $P_s$ | sagittal sinus pressure | mm Hg |
| $G_{[L/R]AM}$ | flow conductance between A/M compartments | (ml/s)/mm Hg |
| $G_{[L/R]MP}$ | flow conductance between M/P compartments | (ml/s)/mm Hg |

**Table 3:** List of diagnostic variables in the six-compartment models

| Symbol | Description | units |
| --- | --- | --- |
| $C_{ic}$ | Intracranial compliance | ml/ mm Hg |
| $r_d$ | Representative vessel radius | cm |
| $Q_{[L/R]AM}$ | distal flow between A/M compartments | ml/s |
| $Q_{[L/R]MP}$ | distal flow between M/P compartments | ml/s |
| $Q_{cAA}$ | anterior distal flow | ml/s |
| $Q_{cPP}$ | posterior distal flow | ml/s |

The most significant quantitative difference between models #2 and #3 for practical nICP estimation is in simulation speed (measured by *r*_3_). Model #1 is an order of magnitude faster than model #2, but both operate considerably faster than real time. Either is suitable for an operational system for clinical support, unlike configuration #3 which is an order of magnitude slower than wall time under the same forcing. Other uni-directional simulations of model #3 with idealized half-sine wave pulsatile ABP inlet conditions showed that very short timesteps (*O*(10^−4^) seconds or shorter) were required to achieve convergence during systole. Longer timesteps (*O*(10^−2^) seconds) sufficed during constant pressure diastolic phase. This suggests that the model requires *O*(10^3^) evaluations and iterative solution steps to the nonlinear system for each simulated cardiac cycle. A previous study ([37]) reported that each cardiac cycle requires 40 seconds within their highly optimized numerical framework. As their implementation used a one-dimensional AN-CoW, *r*_3_ = 0.0225 is a lower bound of the speed score for model #4.

#### 3.1.3 Additional observations and considerations

It is noteworthy to mention that the physiological ICMs of #3 and #4 do not strictly require pulsatile ABP and may instead be forced with mean ABP, which permits larger timesteps during simulation. With pulsatile forcing, they necessarily resolve fine timescales (*cf*. black inset, Fig 5) inherited from the inflow boundary condition and therefore require extensive and inflexible computation time. However, the CA response ODE includes a timescale parameter *t*_*CA*_ which is taken to be 10 seconds by default. These processes are relatively slow and do not respond to inflow changes much shorter than 1 minute (exp(*t*_*CA*_*/*60) ≈ 1.2 minutes). Experiments with model #3 suggest that q1m mean ABP forcing decreases computational overhead considerably (to approximately *r*_3_ = 1.15, slightly faster than real time) by reducing the numerical stiffness of the system. However, doing so comes at a loss of high-frequency nICP fidelity.

Some higher-frequency nICP components, on the other hand, can be resolved by the simple models at the expense of additional computation time by adjusting hyper-parameters. The efficiency score for models #1 and #2 depends on the window length and parameter update interval. Fig 6 shows a portion of a model #2 simulation for Charis patient #6 using raw ABP together with a 30 second window and 1 second update period. The 29-second overlap of the windows corresponds to 30 seconds of model integration for each simulated second. The computational overhead reduces efficiency *r*_3_ to approximately 0.25, but there is considerable gain in nICP fidelity at the high frequencies as well as strongly improved reduced error (*r*_1_) and increased accuracy (*r*_2_). This demonstrates a latent ability of model #2 to estimate higher frequency components of ICP from ABP without the need for accurate ICM parameter inference on the basis of additional data as in model #3.

### 3.2 Simple model experiments with low-frequency inflow data

Models are able to utilize commonly available ABP summary records under appropriate representation. Models #1 and #2 have stricter expectations of data frequency for accurate pulse resolution, but minute-interval timeseries of pressures and heart rate are sufficiently informative to estimate nICP under those models. This is accomplished by broadcasting the q1m data onto a representative waveform with constant diastole, systole, and heart rate between records, and appropriately downsampling the result for use as model forcing. The patient-specific waveform parameters are estimated from a short (5–10 second) interval of high-frequency observed ABP with negligible impact on efficiency *r*_3_.

The nICP estimates based on raw and q1m ABP are difficult to distinguish, which shows that q1m ABP can be used with little impact on accuracy. Fig 7 shows model #2 nICP estimates of the original (blue dashed) and q1m (*γ*_6_, solid blue) solutions for patient #6 compared with the ICP observation (corrected at 243.83 minutes as before). The skills of the model running under accurately reconstructed q1m data are nearly identical to the original; mean error *r*_1_ actually increases by about 3% while there is no qualitative difference in clincal accuracy or efficiency. The nICP changes resulting from q1m forcing are generally limited to the end of the simulation period (220–360 minutes), and the smoother inflow data avoids the inaccurate nICP spike around 175 minutes.

Patient-specific waveform parameters are not necessary for accurate nICP estimation by the simple models, but q1m ABP summaries must include heart rate. The regressive single-compartment model #2 is based only on the peri-systolic changes in pressure and flow. One therefore suspects that the patient-specific parameters *γ* are unnecessary as long as the q1m data are accurate. Two additional simulations with model #2 using incorrect q1m transformations are shown in Fig 7 and confirm that that patient-specific parametric waveforms are unnecessary in these models. The waveforms of Charis patients #8 and #9 clearly differ from those of patient #6 in the post-peak shape, but the associated nICP estimates are qualitatively indistinguishable from the correct one. However, further experiments not detailed here found that model #2 nICP estimates based on q1m ABP without heart rate data were highly inaccurate due to errors in numerical calcuation of the ICM inflow pressure derivative.

### 3.3 Summary of assessments and experiments

#### What is gained and lost from bi-directional coupling

The comparison of models suggests that bi-directional coupling is necessary to accurately track ICP trends when using the simple ICM. Inclusion of this feedback mechanism is crucial in a clinical setting because the bi-directional form of the model is 10% more accurate in correctly specifying critical ICP. The simple ICM, however, is still limited by lack of resolution endogenous IC processes; nICP changes still require corresponding changes in the applied ABP inflow. Interactive coupling between the AN-CoW components and IC model makes the model more prone to potential instabilities during spinup from rest and during periods of noisy ABP inflow data. It is also accompanied by an order of magnitude increase in computation time even when run with non-overlapping analysis windows. However, the coupled simple model in this case is still an order of magnitude faster than clock time. It has sufficient headroom to accommodate a physiologically-representative IC model of (slightly) more complexity (*e*.*g*. [7, 35]) and still retain this desirable advantage. The simple model is able to use short update steps and overlapping analysis windows to estimate higher frequency (∼*O*(10^−2^) Hz) details of ICP at the expense of additional computational time. Model configuration #2 is also able to act on q1m summary ABP data without additional patient-specific parameters to produce nICP estimates nearly-equivalent to those generated from raw ABP data.

#### What is gained and lost from increasing ICM fidelity

Resolving multiple interconnected IC compartments and CA feedback mechanisms offers physiological fidelity at the expense of model complexity. Consequently, the nonlinear ICM has many parameters which could not be accurately specified from available data without advanced inference machinery. Poor identifiability (*i*.*e*. dynamically balanced and representative parameters) of the system required *ad hoc* nICP adjustment to obtain realistic results, and may be improved by parameter tuning. The variance-scaled nICP matches the observed ICP qualitatively but under-performs the simpler models when the ∼4 minute lag in the response signal is accounted for. Simulating the nonlinear ICM toward a solution under pulsatile APB inflow requires many iterations and is considerably slower than real time even with uni-directional coupling. This type of simulation resolves pulse-scale waves within the nICP pulse waves but, as previously noted by [37], they have unrealistically low amplitude. While this type of resolution is valuable for diagnosing changes of IC hemodynamics and CA function [22, 6, 8], it is too computationally expensive for simulations of multiple hours. The computational overhead decreases significantly with smoother, lower-frequency inflow such as mean ABP where high frequency waves are not resolved. Simulations of this type are slightly faster than real time, but are still precluded by the need for ICM parameter estimation. Further exploration of model #3 is needed to evaluate the clinical diagnostic value of simulations driven by non-pulsatile mean ABP over multi-hour periods.

## 4 Discussion

This study presents a multi-component modeling approach to non-invasive estimation of intracranial pressure in the context of multi-hour timescales relevant for producing actionable clinical information. The purpose was to establish a most suitable direction in developing an nICP estimation program that could be applied to commonly available data but fast enough to allow inference. The choices were inclusion of interactive coupling between components or use of a more complex ICM. The key result is that bi-directional coupling of the simple model is sufficiently fast and accurate in test cases and could be implemented using commonly available q1-minute ABP data. The first model component, the AN-CoW, uses long-established electrical analog representations to numerically solve pressure-driven resisted flow through compliant vessels. It represented hemodynamics from the aortic inflow boundary to the ICM interface at CoW termini. The second component, the ICM, models IC processes to obtain non-invasive ICP estimates. Two models differing in complexity and mechanistic fidelity were considered for comparison. The AN-CoW and ICM components could be coupled uni- or bi-directionally, yielding four possible model configurations. We did not implement a highly complex fully-coupled model due to its high computational expense and developmental challenges.

To establish whether bi-directional coupling or increased ICM complexity was more advantageous, model simulation assessed nICP estimation based on accuracy, precision, and speed. A case study involving ABP-based nICP estimation for Charis patient #6 during a slow ICH event revealed that the bi-directionally coupled simple model outperformed both complex and simple uni-directional models. This comparison included both the ability to track the trend of ICP as well as the identification of critical ICP (defined here has ICP*>* 20 mm Hg). Bi-directional coupling to the arterial inflow model was necessary for the simple model (#2) to track the longer-term trend of increasing ICP, but greatly increased computational time compared to its uni-directionally version (#1). In both coupling setups, the simple models were faster than real-time whereas the uni-directional complex model (#3) took nearly six hours to perform a one hour simulation.

The stronger-performing simple model approach may also used on lower-resolution q1m ABP summary data with no patient-specific parametrization of the inflow waveform. Estimates of nICP from the simple bi-directional model (#2) using waveform representations of q1m ABP were approximately equal raw 50 Hz ABP. Results showed that nICP depended only on accurate representation of systolic and diastolic pressures and heart rate; the post-systole pulse shape did not matter. This suggests that coarse clinical q1m data are sufficient drivers for nICP estimation in model #2 either via waveform downsampling during the preprocessing stage or as an aortic model component.

Finally, the computational burden for complex model #3 may be relaxed under mean ABP forcing, but the need for ICM parameter identification limits its utility. Model #3 was also difficult to initialize due to strong parameter dependence. Computation time was an order of magnitude slower than clock time when forced with pulsatile ABP. Simulation speed improved under piece-wise-constant mean (non-pulsatile) ABP forcing, but was only slightly faster than real time. Further, some its many ICM parameters may not be stationary over multi-hour timescales and could require dynamical estimation. This may explain the difficulty in maintaining non-divergent behavior beyond the first hour of simulation. The physiological fidelity offers a potential wealth of clinically useful diagnostic information in the form of internal dynamical parameters and clearly should be used in retrospective applications where nICP waveform features may be desirable. However, the model provides potentially relevant diagnostic information even under mean ABP forcing and this data may be more easily accessible than full ABP series or joint q1m summaries of diastole, systole, and heart rate time series.

The simple models, which rely on statistical analysis of flow and pressure across the middle cerebral artery, are limited by strong stationarity assumptions and are also sensitive to noise in ABP inflow data. The clinical value of model #2 becomes more apparent at longer timescales and is 10% better at correctly predicting whether ICP exceeds 20 mm Hg than model #1. The results were robust under application of a 20Hz low-pass filter to remove noise from raw aortic ABP forcing, as relatively little noise was present in data for patient #6. However, other simulations not shown here suggest that models #1 and #2 are strongly sensitive to noise in aortic signals driving the system, with the latter prone to feedback-driven instabilities as a result of coupling. For example, the errant spike in model #2 nICP simulation around 175 mins (Fig 5, blue line) results from a brief mis-identification of MCA pressure maxima and minima resulting from noisy aortic inflow signals. This effect also presents itself in the model #1 solution (same figure, magenta line). The disruption is brief in the absence of interactive coupling, but poor estimates of IC restance *R* and compliance *C* during the bi-directional simulations inform the following estimation period causing persistent errors during subsequent estimation windows. Shorter update periods increase the potential for instability and linearly increases computational time (*i*.*e*. decreases *r*_3_). Use of longer analysis windows to increase the signal-to-noise ratio risks violating the stationarity assumptions for IC resistance, compliance, and ICP in the single-compartment configuration. The waveform representation of q1m data is sufficiently smooth to avoid many of these problems; the spurious feature around 277 minutes in the *γ*_3_-simulation of Fig 7 remains undiagnosed.

The main results of this work are summarized here:

1. Inclusion of feedback between ICM and AN-CoW components improves tracking of higher order trends over multi-hour timescales. The bidirectionally-coupled single-compartment model #2 features more accurate resolution of low-frequency ICP components than the uni-directionally coupled model.
2. The nICP estimates using q1m ABP data projected onto pulsatile waveforms are qualitatively similar to those obtained using high-frequency APB data. However, q1m summary data must include heart rate in addition to diastolic and systolic pressures.
3. Patient-specific waveforms are *not* required to use q1m ABP as simple model inflow data; the quality of nICP depends neither numerically nor empirically on resolving post-systole components of patient waveforms.
4. Model #2 has stronger potential for multi-hour applications since no parameters are required, can be run using commonly available data, and runs about seven times faster than real time.
5. The large number of parameters within the complex, nonlinear ICM of model #3 suffers from difficult identifiability, and poorly-specified parameters led to divergent or unrealistic behavior. It could not be adequately configured from available data for stable, multi-hour simulations.
6. The temporal resolution of model #3 is inherited from the aortic inflow. Under pulsatile APB forcing, it resolves nICP waveforms but requires extensive computation time. If properly resolved in amplitude by a well-configured and identified model, such waveforms could be used to characterize autoregulatory and adaptive capacity.
7. Near operational nICP estimation speed is plausible with model #3 under forcing by low-frequency or strongly-smoothed ABP, but not pulsatile ABP. Without additional time and data for individualized ICM parameter inference, clinical application may be limited.

### 4.1 Overcoming model limitations: refinement and assimilation

The simple bi-directional model (#2) is a strong candidate to build upon, but it has limitations. Both models #1 and #2 failed to accurately track the ICP trend and variablity of patients suffering intracranial hemorrhage or stroke. The presence of raw ABP noise and large waveform variance may play confounding roles in this limitation. It may also be that idealized physiology limits simple model applicability, as changes to underlying physiology during more acute brain injuries are not representned in the ICM derivation. The simple models base their nICP estimates only on nonlinear and statistical relationships between aortic inflow data and resolved/estimated physiology. They also do not account for many important aspects crucial to clinical decision making process including patient age; injury mechanism; imaging findings; or treatments such as sedation, neuromuscular blockade, osmolar therapy, and ventilation strategy.

Further investigation is warranted to assess whether simple models can account for changes in ICP dynamics arising from unresolved intracranial processes. The MCA pressure and flow used are determined largely (or entirely in uni-directional case) from pulsatile aortic inflow, rather than from systemic ABP and localized CBF data streams as the formulation of [16] intends. Statistical estimation of bulk physiological parameters (*viz*. IC resistance *R* and compliance *C* of models #1 and #2) may not appropriately reflect diminished or exhausted intracranial adaptive capacity. This drawback may manifest itself as inaccurate nICP estimates for cases where ICM dynamics are more sensitive to changes in those parameters or where the physiological range of those parameters changes. For example, models #1 and #2 do not impose upper bounds on IC compliance to reflect thresholds of CA processes or other exhausted adaptability. Additional joint ABP/ICP clinical data are forthcoming and will allow us to more precisely identify the domain of applicability for this model.

Overcoming model #2 limitations to estimate nICP for some patients may require a more complex ICM or inclusion of additional dynamically-controlled parameters. Many patients of clinical concern have more complicated injuries including intracranial hemorrhages and strokes, like the other patients in Charis dataset. The recorded ABP-ICP timeseries of these patients include periods of complex and more rapidly evolving ICP regimes. One such critical ICH period is evident for patient #5, a 21 year-old female with TBI and subdural hematoma, whose ICP rises concerningly from 21 mm Hg to 29 mm Hg over a 47 minute period (Fig 8, red line). The median amplitude within the observed ICP q1m mean is about 0.47 mm Hg relative to its ∼11 minute moving average, which is four times larger than in the record of patient #6 presented previously (*c*.*f*. **Selected Charis patient joint ABP-ICP timeseries**. Fifty Hz ABP (left column) and associated ICP (right column) for Charis patients #1, #5, #6, #8, and #9 (rows top to bottom, respectively, labeled at left). Vertical axes corresponds to pressure measurements in mm Hg and horizontal axes show time in hours. Blue regions outline the raw pulsatile signal, with the red curve identifying the signal smoothed over 2 minute window by a cubic polynomial filter to illustrate the scale of local signal variability. Patient #6 was chosen for benchmarking on the basis of low noise in joint ABP-ICP signals and interpretability of ICP dynamics over a several-hour time period. Patient #5 was chosen for idealized optimizability experiments due the timescale of ICP dynamics continuity over a multi-hour interval, and the decorrelation between ABP and ICP during this period. Patients #8 and #9 are included here since their ABP waveform shapes strongly contrast those of patient #5 (*cf*. Figure 7 inset)). Similar variability exists even in the q1m average ABP inflow signal, and may confound model performance. The quality of nICP estimates for such cases benefits from well-optimized optimization model components, but may require additional machinery to drive model dynamics beyond its inherent ability. Two possible directions of ongoing research are motivated within the modeling framework presented.

#### Increased sophistication

A simple model of increased complexity may account for changes in ICP that arise from IC mechanisms, widening the applicability of the framework of model #2. To broaden the scope of potentially modelable cases, other lumped parameter ICMs which offer both increased physiological fidelity and low-computational overhead may be considered. In particular, two simple models which offer increased IC process resolution and relevant internal parametrization for more detailed clinical diagnostics for CA function assessment have been presented by [35] and [7]. Both are directly representable within the electrical analog framework electrical circuit forms are presented in [11] and contain internal CA mechanisms. Either may easily fit bi-directionally within the existing framework as alternate ICM components with sufficiently fast algorithms for the predictive desire discussed above. These models (and variations thereof utilizing the statistical simplification of [16]) are part of continuing development within the general purview of this research.

#### Additional Parametrization

Another method of applying the simple model to complex cases involves augmented boundary control as a proxy for unresolved processes within a statistical parameter estimation scheme. While patient-specific optimization is beyond the scope of this study, additional experiments applying the model to ABP-ICP timeseries of interest show that model #2 is sufficiently robust to track ICP throughout these complex regimes. This requires the addition of optimized scaling parameters and non-stationary modulation of the relationship between ABP inflow at the aorta and the ICM inflow from the middle cerebral artery. Fig 8 compares observed ICP to nICP estimates obtained from simulations employing independent modulation parameters in addition to partially optimized scaling. Here, the optimized scaling parameters are (*θ*_*l*_, *θ*_*r*_, *ω*_*l*_, *ω*_*r*_, *R*_*term*_) = (0.8, 1, 0.84, 0.93, 1). Simulations shown in the figure use a low-frequency non-stationary gain parameter to vary ABP inflow within the model. Specifically, the inflow source is adjusted by a gain parameter *G*:ABP(*t*) ← ABP(*t*) · (1 + *G*(*t*)). The gain is specified as a piece-wise linear function at regular intervals with |*G*(*t*) | ≤ 0.15. This low-dimensional representation of *G* is effectively determined by a set of discrete parameters, which are potentially estimable from other patient record data. Determining values of gain *G* would involve placing the current ABP-to-nICP model within the framework of a data assimilation system, which provides a meaningful way of automatically constraining uncertainties due to inaccurate parameters and unresolved physiology. Such systems require extensive computational overhead, although some methods such as the empirical Kalman-type methods can take advantage of parallelization to maintain quasi-operational estimation provided that the underlying nICP estimation model is sufficiently fast. rheological parameters deoxyribonucleic acid Fig 8 illustrates the potential of this approach by including two additional model #2 simulations (magenta and cyan curves) using 8 and 24 equally-spaced control points to linearly vary ABP inflow signal. In the absence of this additional control, the scale-parameter optimized model (blue curve) fails to track major waves in observed ICP and is likely of little clinical value. The simulation using 8 additional parameters is more dynamic and resolves a portion of the hypertensive event over 130–150 minutes, but misses its onset and underestimates peak pressure by over 1.5 mm Hg. The mean error *r*_1_ decreases from 8.75 to 6.85 using this additional control. The simulation using 24 additional parameters improves nICP estimation further, qualitatively matching the rising trend of observed ICP from 100–150 minutes as well as the quantitative value of the maximum pressure. The *r*_1_ score is 5.6 under this stronger control, although much of the error is attributed to the consistent positive bias of about 0.7 mm Hg during the final hour of simulation. Either of these solutions is expected to be clinically useful as they indicate the presence of these dynamics which could not be resolved by the model from ABP alone. This example suggests that the model is capable of reproducing observed ICP from ABP for complex cases with additional non-stationary parametric optimization, and motivates ongoing work in that direction. Practical applications require estimation of these additional parameters whereas they were specified *a priori* in this illustration. Nevertheless, this example further underscores the need for simple, fast models to meet the goal of providing timely, relevant nICP estimation over multi-hour timescales.

### 4.2 Forecast potential for clinical support

A model-based forecast system based on bi-bidirectionally coupled model #2 can potentially inform clinicians of possible impending problems by extrapolating parameter and dynamical trends into the future. Such a system would greatly benefit both clinical decision support and care-level logistics by indicating possible changes in patient status with sufficient lead time to adjust room, equipment, and staff. This may also give practitioners advance warning with a time frame for planning treatments, permitting earlier and lower-risk interventions to combat IC hypertension. Recent works [1, 31, 33, 40] include machine-learning approaches to ABP prediction, and could be used in conjunction with the presented methods for short-term prediction of nICP. The application of these algorithms to low sample-rate q1m ABP records has not been reported in the literature.

The speed of model #2 indicates it is a plausible candidate for use within a statistical estimation and forecast scheme which requires many forward model integrations. The accurately identified parameters together with acceptable simulation speed adds the possibility of practical forecast capabilities on the basis of trends in diagnostically computed model parameters. For the applications discussed in this work, distributional trends and higher order moments in ICM resistance and compliance may be inferred from robustly optimized model #2 simulations of a patient’s relevant history. This statistical information may then be utilized to predict possible future ICP outcomes under current ABP measurements or ABP forecast, potentially providing valuable and timely clinical decision support for caretakers and facility management.

### 4.3 Ongoing work

The original hypothesis was that the high degree of physiological fidelity of the complex multi-compartment model would provide the most diagnostic information from available data. It also had numerous model parameters which could be inferred from patient data in the longer view of the research program, which is to aid in patient-specific clinical support. This work began with an attempted implementation of model #4 using a spatially-resolved vascular system as in [29], which had recently been used within a data assimilation system [37]. We adopted the 0D AN-CoW to eliminate complicated propagation of waves across AN-CoW bifurcations and to increase computational speed. This choice did not easily allow for bi-directional coupling with the nonlinear ICM. These difficulties with bi-directional coupling arose from enforcing CoW-ICM pressure and flow agreement during the iterative solution of the complex ICM. MCA boundary outflow of the AN-CoW was saved for offline development of model #1 and for offline debugging of the stand-alone ICM of model #4, which became model #3. However, solving the complex non-linear ICM system by iterative means was slow, and difficult to initialize from available data due to strong parameter dependence. The simple model #1 was easily coupled bi-directionally to the AN-CoW system, becoming model #2. This incorporated model was found to better track lower-frequency trends in ICP and could resolve higher-frequency ICP waves with additional computational cost. The lack of sophistication and parametrization within the simpler schemes motivates the need for additional external parameters to control the solution in more complex patient cases. This work describes these developments and informs possible directions of ongoing research toward the ultimate goal of developing operational nICP estimation suitable for clinical use.

The need for additional inference is clear for application of models #2 and #3, but there are substantive differences in methodology and potential benefits. The simple model #2 lacks parameters and therefore requires no additional data, but is limited to applications where there is a strong correlation between systemic ABP and ICP response. To overcome that limitation, use of a more sophisticated ICM or augmentation with external parametric control over ABP inflow are proposed above (subsection 4.1). On the other hand, model #3 has numerous internal parameters which need to be properly inferred for meaningful simulations, but has the advantage of very strong parameter interpretability. Lack of physical meaning for proposed control parameters of model #2 limits the quality of information an optimized solution may produce beyond improving nICP estimation, while the investment of time to infer parameters in model #3 yields a wealth of clinically relevant knowledge. Furthermore, patient-specific parameters in the ICM of model #3 are numerous and interacting, but are also presumed to be stationary and thus potentially inferable from historical data using traditional methods (*e*.*g*. MCMC estimation or optimization). In contrast, the control mechanism proposed for model #2 requires only a few parameters, but they are distributed in time with unknown temporal correlations. A plausible method of estimation in this situation is via ensemble filtering, but the necessary mapping between typical clinical data and the control parameters is currently unknown and requires further development. Given that estimation of nICP is the primary objective of this project and inference requires many repeated simulations, continued development of inference machinery for model #2 is likely the best choice.

The long term vision of this project remains the development of a bi-directionally coupled model with anatomical fidelity (*i*.*e*. model #4) fast enough for pre-emptive diagnostic use. One path toward this goal is a hybridization of methodologies that integrates an ICM of intermediate complexity under piece-wise stationarity assumptions akin to those of the simple models. Possible ICMs include those mentioned previously and a simplified (*e*.*g*. linearized) counterpart of model #3. This should reduce the burden of computational time of the complex model and allow it to be more easily coupled interactively to the upstream vascular component. Such a model would further benefit from highly interpretable inference based on data available when administering care, with the additional advantage of supporting summary ABP inflow. A remaining question is whether a model formulated in this way can be made fast enough to provide timely and clinically actionable information.

## [SI-1]Detailed Model Descriptions

### 4.4 Zero-dimensional vessel parametrization

Physical parametrization of vessel-level hemodynamics in the AN-CoW involves vessel dimensions (cross-sectional area *A*_0_ and length *l*), material properties (vessel linear compliance *∂P/∂A*, vessel elasticity constant *β*, and blood density *ρ*), and a friction scaling term (*χ*, which depends on vessel mechanical properties and flow profile [14]). A local elastic pressure model *P* = *P* (*A*; *β, A*_0_) is adopted from [24], where

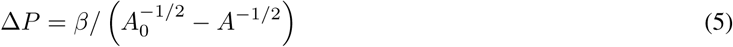

where Δ*P* is the change in internal pressure with respect to transmural vessel pressure. Parameters defining the passive electrical components of each vessels are resistance *R*, capacitance *C*, and inductance *Z*. These may be define approximated [23] from the physical parameters according to

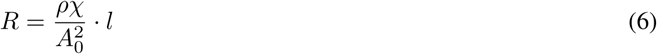

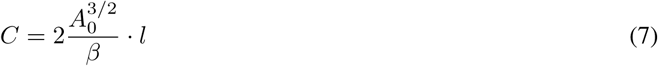

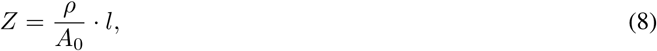

respectively. The relationship between *A* and vessel radius *r* is elementary.

### 4.5 Six-Compartment ICM details

The six-compartment model [29] is computationally centered at the the distal cerebral arterial bed represented by the complaint structures *C*_*d*$*X*_) of each of the 6 territories visible in Figure 2. The physiological model combines Laplace’s for law balancing wall tension with arterial pressure. Representations for tension as functions of pressure *P* gives:

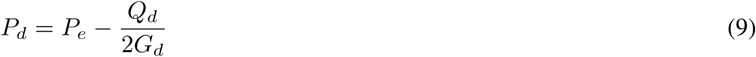

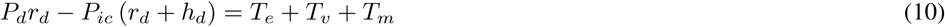

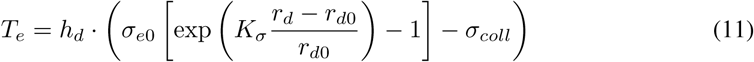

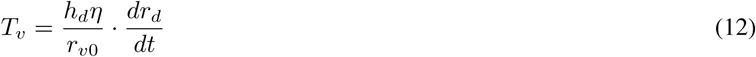

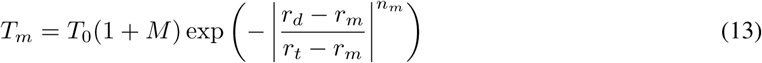

where subscripts *e, d* correspond to the proximal and distal arterial beds. Values of *P*_*e*_ correspond to interface pressure between the vascular network and the ICM at CoW outflows, whose character varies depending on the coupling method. Values of *Q*_*d*_ and Δ*Q*_*coll*_ represent flows determined by transported fluid balances of each cerebral compartment.

The tension term *T*_*m*_ models cranial auto-regulation (CA) through modulation of the state-dependent variable *M* ∈ [−1, 1] determining vaso-dilation/constriction of effective vascular radius *r*_*d*_ of each compartment. CA is modeled by the dynamics of a feedback mechanism *ξ* that aims to relax the distal lumped flow *Q*_*d*_ to a target flow *Q*_*n*_ over timescale *t*_*CA*_ with gain factor; the adjustment ODE determines *M* as:

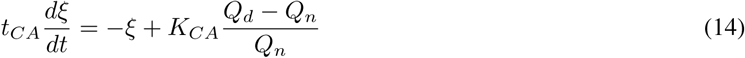

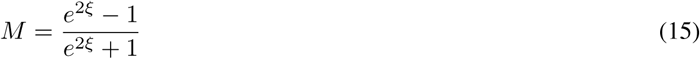

The volume balance for each territory is given in terms of its effective vascular radius *r*_*d*_:

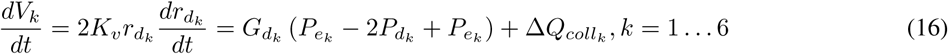

where 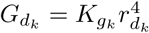.

The collateral flow volumes 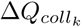 are determined by pressure-difference-driven flows between adjacent compartments (see Eqns.25[29]).

Once blood flow distribution of each compartment is represented, the common ICP value *P*_*ic*_ for the component is the solution to the differential equation

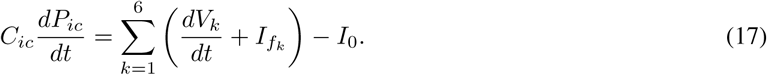

However, the undetermined ICP influences both CSF outflow *I*_*o*_ and the bed-wise CSF production rates 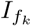 as well as the the intracranial compliance *C*_*ic*_. Equation (17) must therefore be solved with the nonlinear terms (for *k* = 1 … 6)

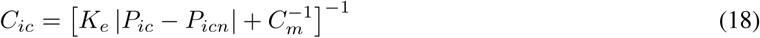

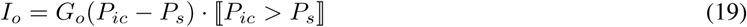

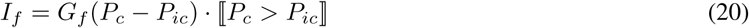

with double brackets denoting test operators.

#### 4.5.1 Numerical Implementation

The system is represented numerically as

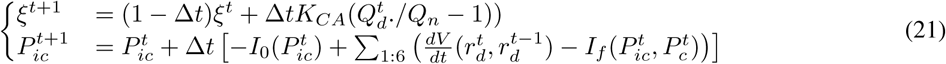

and solved by minimizing the nonlinear function *R*(*x*) = |*M* (*x*)*x* − *b*(*x*)| where

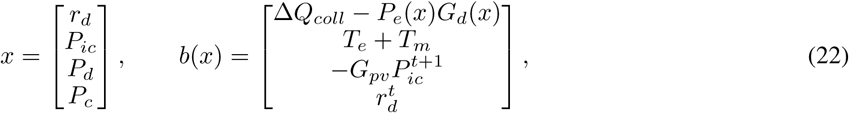

and

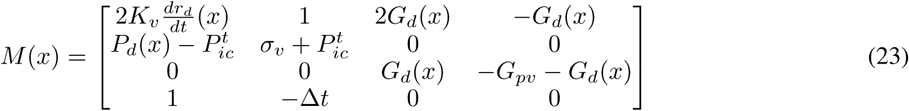

and is initialized using known or computed values at *t*.

The entry-wise values of the optimum *x* provide updated values of its constituents at time *t* + 1. The last row of the system enforces a finite difference approximation to *dr/dt*, but it makes the system explicitly Δ*t*-dependent.

### 4.6 Single-compartment

The single compartment model[16] seeks to identify IC compliance and resistance by regressing features of the forcing waveforms across temporal intervals. In particular, IC inflow *Q* and *P* are represented here in electrical analog form (see Figure 2. The principal assumptions are that ICP, IC resistance *C*, and IC resistance *R* are constant over the regression interval, in which case the system reduces to an RC circuit with governing equation

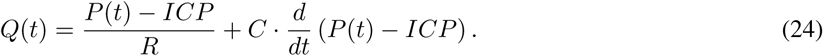

During each systolic upswing {*t*_*a*_ <= *t* <= *t*_*b*_}, flow through the resistance is assumed to be small and the entire flow is stored compliantly; therefore, the value of *C* is estimated by regressing the net inflow volume against the change in pressure during that interval:

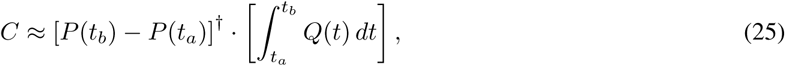

with (·)^†^ indicating the pseudo-inverse/least-squares matrix. Identification of intervals *{*[*t*_*a*_, *t*_*b*_]} proceeds by identifying roughly the times of minimum and maximum applied pressure.

With the ICM inflow decomposed into resistive and capacitive flows, the former flow is calculated using the estimate of *C*

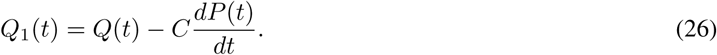

ICP, assumed to be constant over the interval, is the difference between applied pressure *P* (*t*) and the pressure lost to forcing resisted flow:

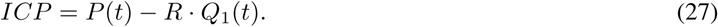

Evaluating at pairs of nearby times *t*_1_, *t*_2_ eliminates *ICP*, and the value of *R* is determined by regressing the change in pressure against the corresponding change in resistive flow:

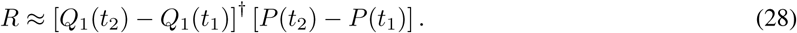

Model #2 uses estimated values of *R* and *C* to estimate nICP directly via Equation (27)[16], whereas model #3 updates *R* and *C* and simulates nICP.

#### SI-2

Additional Figs, etc

## References

[1] Robabeh Abbasi, Mohammad Hassan Moradi, and Seyyedeh Fatemeh Molaeezadeh. Long-term prediction of blood pressure time series using multiple fuzzy functions. In 2014 21th Iranian Conference on Biomedical Engineering (ICBME), pages 124–127. IEEE, 2014.

[2] Charles A Adams, Deborah M Stein, Jonathan J Morrison, and Thomas M Scalea. Does intracranial pressure management hurt more than it helps in traumatic brain injury? Trauma surgery & acute care open, 3(1), 2018.

[3] William M Armstead. Cerebral blood flow autoregulation and dysautoregulation. Anesthesiology clinics, 34(3):465–477, 2016.

[4] Marcella Balestreri, Marek Czosnyka, Peter Hutchinson, Luzius A Steiner, Magda Hiler, Piotr Smielewski, and John D Pickard. Impact of intracranial pressure and cerebral perfusion pressure on severe disability and mortality after head injury. Neurocritical care, 4(1):8–13, 2006.

[5] Chen-Huan Chen, Erez Nevo, Barry Fetics, Peter H Pak, Frank CP Yin, W Lowell Maughan, and David A Kass. Estimation of central aortic pressure waveform by mathematical transformation of radial tonometry pressure: validation of generalized transfer function. Circulation, 95(7):1827–1836, 1997.

[6] Marek Czosnyka and John D Pickard. Monitoring and interpretation of intracranial pressure. Journal of Neurology, Neurosurgery & Psychiatry, 75(6):813–821, 2004.

[7] Marek Czosnyka, Stefan Piechnik, Hugh K Richards, Peter Kirkpatrick, Piotr Smielewski, and John D Pickard. Contribution of mathematical modelling to the interpretation of bedside tests of cerebrovascular autoregulation. Journal of Neurology, Neurosurgery & Psychiatry, 63(6):721–731, 1997.

[8] Jun-Yu Fan, Catherine Kirkness, Paolo Vicini, Robert Burr, and Pamela Mitchell. Intracranial pressure waveform morphology and intracranial adaptive capacity. American Journal of critical care, 17(6):545–554, 2008.

[9] Andrea Fanelli, Frederick W Vonberg, Kerri L LaRovere, Brian K Walsh, Edward R Smith, Shenandoah Robinson, Robert C Tasker, and Thomas Heldt. Fully automated, real-time, calibration-free, continuous noninvasive estimation of intracranial pressure in children. Journal of Neurosurgery: Pediatrics, 24(5):509–519, 2019.

[10] Ary L Goldberger, Luis AN Amaral, Leon Glass, Jeffrey M Hausdorff, Plamen Ch Ivanov, Roger G Mark, Joseph E Mietus, George B Moody, Chung-Kang Peng, and H Eugene Stanley. Physiobank, physiotoolkit, and physionet: components of a new research resource for complex physiologic signals. circulation, 101(23):e215–e220, 2000.

[11] Christopher Hawthorne and Ian Piper. Monitoring of intracranial pressure in patients with traumatic brain injury. Frontiers in neurology, 5:121, 2014.

[12] Lucy He, Travis R Ladner, Sumit Pruthi, Matthew A Day, Aditi A Desai, Lori C Jordan, and Michael T Froehler. Rule of 5: angiographic diameters of cervicocerebral arteries in children and compatibility with adult neurointerventional devices. Journal of neurointerventional surgery, 8(10):1067–1071, 2016.

[13] Xiao Hu, Valeriy Nenov, Marvin Bergsneider, Thomas C Glenn, Paul Vespa, and Neil Martin. Estimation of hidden state variables of the intracranial system using constrained nonlinear kalman filters. IEEE Transactions on Biomedical Engineering, 54(4):597–610, 2007.

[14] Thomas Jr Hughes and J Lubliner. On the one-dimensional theory of blood flow in the larger vessels. Mathematical Biosciences, 18(1-2):161–170, 1973.

[15] Gerard N Jager, Nico Westerhof, and Abraham Noordergraaf. Oscillatory flow impedance in electrical analog of arterial system: representation of sleeve effect and non-newtonian properties of blood. Circulation research, 16(2):121–133, 1965.

[16] Faisal M Kashif, George C Verghese, Vera Novak, Marek Czosnyka, and Thomas Heldt. Model-based noninvasive estimation of intracranial pressure from cerebral blood flow velocity and arterial pressure. Science translational medicine, 4(129):1–9, 2012.

[17] Marium Naveed Khan, Hussain Shallwani, Muhammad Ulusyar Khan, and Muhammad Shahzad Shamim. Noninvasive monitoring intracranial pressure–a review of available modalities. Surgical neurology international, 8, 2017.

[18] Meeri N Kim, Turgut Durduran, Suzanne Frangos, Brian L Edlow, Erin M Buckley, Heather E Moss, Chao Zhou, Guoqiang Yu, Regine Choe, Eileen Maloney-Wilensky, et al. Noninvasive measurement of cerebral blood flow and blood oxygenation using near-infrared and diffuse correlation spectroscopies in critically brain-injured adults. Neurocritical care, 12(2):173–180, 2010.

[19] Nam Kim, Alex Krasner, Colin Kosinski, Michael Wininger, Maria Qadri, Zachary Kappus, Shabbar Danish, and William Craelius. Trending autoregulatory indices during treatment for traumatic brain injury. Journal of clinical monitoring and computing, 30(6):821–831, 2016.

[20] William D Lakin, Scott A Stevens, Bruce I Tranmer, and Paul L Penar. A whole-body mathematical model for intracranial pressure dynamics. Journal of mathematical biology, 46(4):347–383, 2003.

[21] Niels A Lassen. Cerebral blood flow and oxygen consumption in man. Physiological reviews, 39(2):183–238, 1959.

[22] JJ1 Lemaire, T Khalil, F Cervenansky, G Gindre, JY Boire, JE Bazin, B Irthum, and J Chazal. Slow pressure waves in the cranial enclosure. Acta neurochirurgica, 144(3):243–254, 2002.

[23] Vuk Milišić and Alfio Quarteroni. Analysis of lumped parameter models for blood flow simulations and their relation with 1d models. ESAIM: Mathematical modelling and numerical analysis, 38(4):613–632, 2004.

[24] Mette S Olufsen. Structured tree outflow condition for blood flow in larger systemic arteries. American journal of physiology-Heart and circulatory physiology, 276(1):H257–H268, 1999.

[25] Mette S Olufsen, Charles S Peskin, Won Yong Kim, Erik M Pedersen, Ali Nadim, and Jesper Larsen. Numerical simulation and experimental validation of blood flow in arteries with structured-tree outflow conditions. Annals of biomedical engineering, 28(11):1281–1299, 2000.

[26] Alfredo L Pauca, Michael F O’Rourke, and Neal D Kon. Prospective evaluation of a method for estimating ascending aortic pressure from the radial artery pressure waveform. Hypertension, 38(4):932–937, 2001.

[27] Alfredo L Pauca, Stephen L Wallenhaupt, Neal D Kon, and William Y Tucker. Does radial artery pressure accurately reflect aortic pressure? Chest, 102(4):1193–1198, 1992.

[28] Michael J Rosner, Sheila D Rosner, and Alice H Johnson. Cerebral perfusion pressure: management protocol and clinical results. Journal of neurosurgery, 83(6):949–962, 1995.

[29] Jaiyoung Ryu, Xiao Hu, and Shawn C Shadden. A coupled lumped-parameter and distributed network model for cerebral pulse-wave hemodynamics. Journal of Biomechanical Engineering, 137(10):101009, 2015.

[30] Bernhard Schmidt, Jurgen Klingelh ofer, Jens J urgen Schwarze, Dirk Sander, and Ingo Wittich. Noninvasive prediction of intracranial pressure curves using transcranial doppler ultrasonography and blood pressure curves. Stroke, 28(12):2465–2472, 1997.

[31] Costas Sideris, Haik Kalantarian, Ebrahim Nemati, and Majid Sarrafzadeh. Building continuous arterial blood pressure prediction models using recurrent networks. In 2016 IEEE International Conference on Smart Computing (SMARTCOMP), pages 1–5. IEEE, 2016.

[32] Nino Stocchetti and Andrew IR Maas. Traumatic intracranial hypertension. New England Journal of Medicine, 370(22):2121–2130, 2014.

[33] Peng Su, Xiao-Rong Ding, Yuan-Ting Zhang, Jing Liu, Fen Miao, and Ni Zhao. Long-term blood pressure prediction with deep recurrent neural networks. In 2018 IEEE EMBS International Conference on Biomedical & Health Informatics (BHI), pages 323–328. IEEE, 2018.

[34] Samon Tavakoli, Geoffrey Peitz, William Ares, Shaheryar Hafeez, and Ramesh Grandhi. Complications of invasive intracranial pressure monitoring devices in neurocritical care. Neurosurgical focus, 43(5):E6, 2017.

[35] Mauro Ursino and Carlo Alberto Lodi. A simple mathematical model of the interaction between intracranial pressure and cerebral hemodynamics. Journal of Applied Physiology, 82(4):1256–1269, 1997.

[36] Mauro Ursino and Carlo Alberto Lodi. Interaction among autoregulation, CO2 reactivity, and intracranial pressure: a mathematical model. American Journal of Physiology-Heart and Circulatory Physiology, 274(5):H1715–H1728, 1998.

[37] Jian-Xun Wang, Xiao Hu, and Shawn C Shadden. Data-augmented modeling of intracranial pressure. Annals of Biomedical Engineering, 47(3):714–730, 2019.

[38] Nicolaas Westerhof, Frederik Bosman, Cornelis J De Vries, and Abraham Noordergraaf. Analog studies of the human systemic arterial tree. Journal of biomechanics, 2(2):121–143, 1969.

[39] Mark H Wilson. Monro-Kellie 2.0: The dynamic vascular and venous pathophysiological components of intracranial pressure. Journal of Cerebral Blood Flow & Metabolism, 36(8):1338–1350, 2016.

[40] Bing Zhang, Zhiyao Wei, Jiadong Ren, Yongqiang Cheng, and Zhangqi Zheng. An empirical study on predicting blood pressure using classification and regression trees. IEEE access, 6:21758–21768, 2018.

